# The FKBP42, TWISTED DWARF1, prioritizes auxin over brassinosteroid transport by peptidyl-prolyl *cis-trans* isomerization of ABCB1

**DOI:** 10.64898/2026.07.24.740493

**Authors:** Tashi Tsering, Francesca Iacobini, Jennifer Sapia, Martin Di Donato, Aurélien Bailly, Peter Hunyadi, Xiaoyu Xia, Hong Wei, Tamás Hegedűs, Markus M. Geisler

## Abstract

The ATP-binding cassette (ABC) transporter ABCB1 transports both auxins and the brassinosteroid brassinolide (BL). ABCB1-mediated auxin (IAA) transport depends on interaction with the FKBP42 protein TWISTED DWARF1 (TWD1), yet the mechanism underlying substrate selectivity remains unclear.

Here, we confirm dual IAA and BL transport by ABCB1 and show that the two substrates compete for ABCB1-mediated transport. We demonstrate that a conserved proline residue (P1008) is required for IAA, but not BL, transport. Furthermore, we identify TWD1 as a calmodulin-activated peptidyl-prolyl *cis-trans* isomerase (PPIase) that selectively enhances IAA, but not BL, transport through isomerization of the E1007-P1008 peptide bond. Loss of TWD1 PPIase activity abolishes ABCB1-mediated IAA export without affecting BL transport, revealing a regulatory mechanism that prioritizes auxin over brassinosteroid transport. Cryo-EM, molecular docking, and molecular dynamics simulations of wild-type ABCB1 and the ABCB1^P1008G^ variant suggest that the P1008 loop mediates long-range communication between the nucleotide-binding domain surface and the substrate-binding pocket, thereby providing a structural mechanism by which TWD1 modulates substrate specificity.

In summary, our findings establish the structural basis of ABCB1 substrate selectivity and uncover a post-translational mechanism that selectively regulates auxin transport while preserving brassinosteroid transport.

**One Sentence Summary:** TWD1 prioritizes ABCB1-mediated auxin over brassinosteroid transport

## Introduction

Multidrug resistance (MDR) poses a significant challenge in cancer treatment and the management of pathogenic yeast infections, largely due to the overexpression of ATP-binding cassette (ABC) transporters (Khunweeraphong and Kuchler 2021). These transporters, particularly members of the ABCB subfamily, actively efflux chemotherapeutic agents and antifungal drugs, reducing their intracellular concentrations and efficacy (Khunweeraphong and Kuchler 2021; Modok et al. 2006). Mammalian ABCBs are known for their broad substrate specificity, which is a key factor in their role in MDR. Despite extensive research, the molecular determinants of substrate specificity in ABC transporters remain incompletely understood.

In plants, the ABC family is highly expanded (Verrier et al. 2008; Anfang and Shani 2021; Park et al. 2017; Hwang et al. 2016; Do et al. 2021; Banasiak and Jasinski 2022). A subset of ABCBs in the model plant Arabidopsis and several crop plants plays a critical role in the transport of the major auxin, IAA (indolyl-3-acetic acid), which is a primary determinant of growth and development (Vanneste and Friml 2009; Hammes et al. 2022; Geisler 2021; Geisler et al. 2005; Chen et al. 2023).

Over the last two decades, convincing indirect and direct evidence employing different homologous and heterologous expression and transport systems has been provided that a subclass of ABCBs from the model plant *Arabidopsis thaliana* and several crop plants catalyze the specific transport of auxin (Chen et al. 2023; Ofori et al. 2018; Xu et al. 2014; Kubes et al. 2012; Kamimoto et al. 2012; Yang and Murphy 2009; Titapiwatanakun et al. 2009; Bailly et al. 2008; Bouchard et al. 2006; Geisler et al. 2005). Interestingly, in contrast to mammalian ABCBs, plant ABCBs were shown to own a far higher degree of substrate specificity (Geisler 2024). ABCB1 was shown to transport beside IAA a few auxinic compounds, such as 1-NAA and 2.4-D but not 2-NAA or IBA (Geisler 2021; Blakeslee et al. 2007; Santelia et al. 2005; Geisler et al. 2005). Moreover, root and shoot PAT defects measured on inflorescence stems and seedlings using radiolabeled IAA agree with auxin reporter analyses and developmental read-outs. *Abcb* mutants present defects in auxin-related read-outs, such as epinastic cotyledons, lateral root formation defects, enhanced gravitropism, photomorphogenesis, smaller rosettes (Wu et al. 2007; Noh et al. 2001; Geisler et al. 2005; Chen et al. 2023; Geisler et al. 2003).

A mutagenesis approach has identified E1007-P1008 in ABCB1 to be essential for transport of IAA, which is part of a diagnostic D/E-P motif for auxin-transporting ABCBs, called ATAs (Hao et al. 2020; Geisler and Hegedus 2020). The E-P1008 of ABCB1 is a constituent of the previously mapped contact site between the FKBD of TWD1 and the NBD2 of ABCB1, respectively (Geisler et al. 2003). Based on this signature motif, 11 of the 22 full-size ABCBs of *Arabidopsis thaliana* were suggested to transport auxin, which was recently verified (Tsering et al. 2024).

Recently, Arabidopsis ABCB1 and ABCB19 were shown to transport additionally brassinosteroids (BR), another class of structurally very different growth promoting hormones (Ying et al. 2024; Wei et al. 2025; Chen et al. 2025; Liu and Liao 2025). Cryo-EM structures allowed mapping of BR binding in the hydrophobic cavity of the transmembrane domains (TMDs) via both hydrophobic interactions and hydrogen bonds (Ying et al. 2024; Wei et al. 2025; Chen et al. 2025; Liu and Liao 2025). Surprisingly, all BL-bound and unbound forms of ABCB1 and ABCB19 present almost identical structures with only minor structural changes in transmembrane domains upon BR binding (Ying et al. 2024; Wei et al. 2025) An outward conformation of ABCB1 was so far only found in the presence of IAA and an ATP analog (Chen et al. 2025), although IAA was not co-crystalized in the structure, most likely due to its low affinity to ABCB1. This enhanced specificity of plant ABCBs for a few substrates (here: auxins and brassinosteroids), was recently referred to as substrate multi-specificity (Geisler 2024) and might highlight evolutionary adaptations in substrate recognition compared to their counterparts in animals and fungi (Geisler 2024; Bailly et al. 2011). The mode of action how ABCB1 differentiate and transport these two structurally very distinct substrates is currently unclear (Chen et al. 2025).

Current work has provided evidence that the immunophilin-like FKBP42, TWISTED DWARF1 (TWD1), functions together with HEATSHOCK PROTEIN90 (HSP90) as chaperones of subset of ABCBs (Geisler et al. 2003; Geisler and Hegedus 2020; Geisler et al. 2016; Geisler and Bailly 2007; Tsering et al. 2024). In the Arabidopsis *twd1* mutant, plasma membrane-localized ABCB1,4,19 but no other ATAs are retained on the ER suggesting defects in early ABCB biogenesis (Wang et al. 2013; Wu et al. 2010; Tsering et al. 2024). HSP90 differentially stabilizes ABCB1,4,19 auxin transporters on the plasma membrane in an action that is dependent on TWD1 (Tsering et al. 2024). In agreement, *twd1* and *abcb1,19* mutants show widely overlapping dwarf phenotypes and disoriented (“twisted”) cellular and organ growth (Geisler and Bailly 2007; Geisler et al. 2003; Wang et al. 2013; Wu et al. 2010).

In analogy, the mammalian ortholog of FKBP42/TWD1, FKBP38, associates with and controls steady-state levels of ABCC7/CYSTIC FIBROSIS TRANSMEMBRANE CONDUCTANCE REGULATOR (CFTR) (Aryal et al. 2015; Banasavadi-Siddegowda et al. 2011; Geisler and Hegedus 2020). CFTR is a C-/MRP-type ABC transporter, functioning as a chloride/bicarbonate channel whose mutation is responsible for the genetic disease cystic fibrosis (Cant et al. 2014; Geisler and Hegedus 2020; Csanady et al. 2019). Interestingly, the peptidyl-prolyl *cis-trans* isomerase (PPIase) function of FKBP38 plays an important role in membrane protein biogenesis on the cytoplasmic side of the ER membrane (Edlich et al. 2007a). All the above is suggesting a common, cross-kingdom regulatory mechanism between ABCB1,4,19/TWD1 and CFTR/FKBP38 modules. In those TWD1/FKBP38 would regulate ABCB/CFTR biogenesis correlating with the tetratricopeptide (TPR) repeat domain of TWD1 and FKBP38, respectively, providing the surface for HSP90 interaction (Tsering et al. 2024; Edlich et al. 2007a). In contrast, regulation of transport activity is thought to be associated with the FKBD of TWD1/FKBP38 harboring the PPIase activity (Geisler and Hegedus 2020).

A recent systematic analysis has uncovered a correlation between TWD1 interaction and ABCB auxin transport regulation. Co-expression of ABCB1,4,19 with TWD1 enhanced ABCB-mediated IAA export, while other ATAs, like ABCB6 or ABCB16, were not affected, However, all attempts to validate a PPIase activity on TWD1 have failed (Kamphausen et al. 2002) leaving the mechanistic impact of the conserved P1008 on ABCB1 transport an open question (Geisler and Bailly 2007; Geisler et al. 2003; Wang et al. 2013; Wu et al. 2010).

## Results

In this study, we address the impact of the D/E-P motif on ABCB1-mediated transport of auxin and BR and show that TWD1 by means of an inducible PPIase activity prioritizes auxin over BR transport. In a first step and in order to independently verify previously reported BR transport activities for ABCB1 and ABCB19 (Ying et al. 2024; Wei et al. 2025; Chen et al. 2025), we analyzed efflux of radiolabeled IAA and the BR, brassinolid (BL), from isolated protoplasts prepared from established Arabidopsis *abcb* and *twd1* lines (Bouchard et al. 2006; Geisler et al. 2005; Geisler et al. 2003). IAA (Bouchard et al. 2006; Geisler et al. 2003) and BL showed a similar, gradually reduced efflux behavior for *abcb* single, *abcb* double mutants and *twd1* (Fig. 1a-b), while efflux of the diffusion control, benzoic acid (BA), was not different from Wt (Suppl. Fig. 1a).

**Fig. 1:**
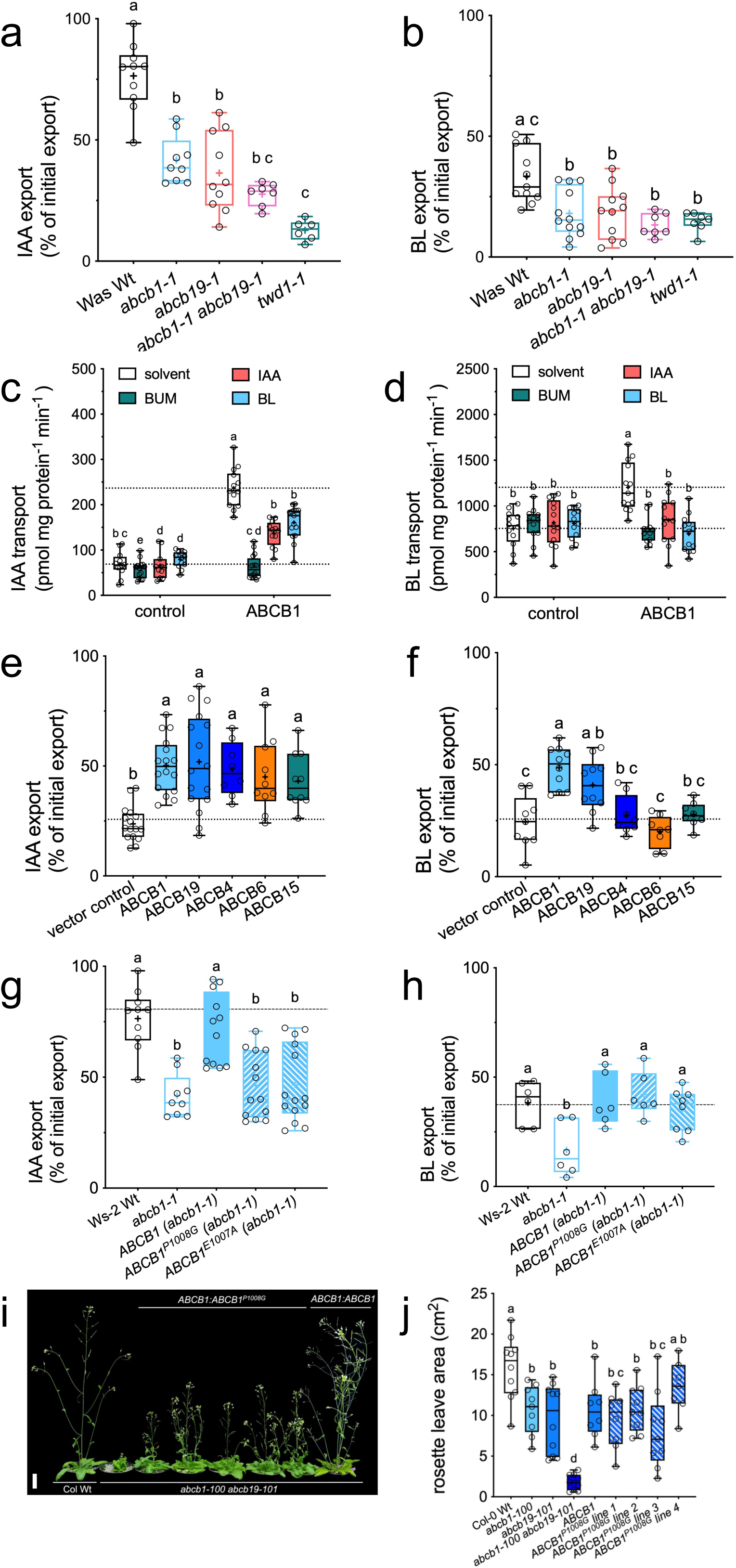
ABCB1 and ABCB19 own a dual IAA and BL transport activity, with IAA transport being dependent on a conserved D/E-P motif. **a-b.** IAA (**a**) and BL (**b**) export from Arabidopsis Wt, *abcb1, abcb19*, *abcb1 abcb19* and *twd1* protoplasts. Significant differences (*p* < 0.05) of means (indicated by “+”) ± SE (n ≥ 8 independent protoplast preparations) were determined using Ordinary One-way ANOVA (Tukey’s multiple comparison test) and are indicated by different lowercase letters. **c-d.** Competition assays of ^3^H-IAA (**c**) and ^3^H-BL uptake (**d**) into tobacco vesicles prepared from vector control (control) or 35S:ABCB1 (ABCB1) transfected leaves. Uptake of 1 nM ^3^H-IAA and ^3^H-BL was measured in the absence (solvent) or presence of 1000x access (1 μM) of non-labelled IAA or BL or in the presence of 50 μM BUM. Significant differences (*p* < 0.05) of means ± SE (n = 3 independent vesicle preparations) were determined using Ordinary one-way ANOVA (Tukey’s multiple comparison test) and are indicated by different lowercase letters. **e-f**. IAA (**a**) and BL (**b**) export from *N. benthamiana* protoplasts after transfection with indicated ABCBs. Significant differences (*p* < 0.05) of means ± SE (n ≥ 8 independent protoplast preparations) were determined using Ordinary One-way ANOVA (Tukey’s multiple comparison test) and are indicated by different lowercase letters. **g-h.** IAA (**g**) and BL (**h**) export from Arabidopsis protoplasts prepared from *abcb1* complemented with Wt (ABCB1) or mutated ABCB1 (ABCB1^P1008G^, ABCB1E^1007A^). Significant differences (*p* < 0.05) of means (indicated by “+”) ± SE (n ≥ 8 independent protoplast preparations) were determined using Ordinary One-way ANOVA (Tukey’s multiple comparison test) and are indicated by different lowercase letters. **i-j.** An auxin transport-incompetent (D1008) version of ABCB1 complements nearly entirely shoot growth defects of *abcb1 abcb19*. Phenotype (**i**) and rosette leaf surface of of 30 dag pot-grown plants (**h**); bar, 2 cm.

Next, we investigated the competitive behavior of both substrates by measuring ATP-dependent uptake into microsomes prepared from tobacco leaves transfected either with ABCB1 or the vector control. To validate our transport system, we employed the ABCB-specific auxin transport inhibitor BUM (Kim et al. 2010) that was recently reported by cryo-EM to bind to a hydrophobic pocket harboring two BUM molecules, which closely overlaps with the BL substrate binding domain (Liu and Liao 2025). As expected, BUM inhibited likewise IAA and BL transport by ABCB1 to vector control level, while its impact on vector control was only marginal (Fig. 1c-d). Interestingly, both IAA and BL (added at a 1000-fold access) were able to significantly compete for both ^3^H-IAA and ^3^H-BL uptake, respectively, while vector control uptake was widely unaffected (Fig. 1c-d).

Subsequently, we tested selected Arabidopsis ABCB isoforms for their IAA and BL transport capacities using protoplasts prepared from tobacco leaves agrobacterium-infiltrated with either ABCB or the vector control. Interestingly, all tested ABCBs transported IAA but beside ABCB1 and ABCB19, neither ABCB4, ABCB6 nor ABCB15 as representatives of ABCB4/21 (Kamimoto et al. 2012; Santelia et al. 2005), ABCB6/20 (Zhang et al. 2018) or ABCB15-18,22 clades (Chen et al. 2023; Hao et al. 2020), respectively, were able to transport BL (Fig. 1e-f).

Previous work had shown that expression of Wt but not of E1007-P1008 mutant versions of ABCB1 was able to export IAA upon expression in tobacco (Hao et al. 2020). We therefore analyzed IAA and BL export from *abcb1* mutant alleles complemented with Wt and P1008G and E1007A mutant versions of ABCB1, respectively, both expressed under their native promoters. As expected, transgenic lines had no obvious phenotype and plant height of ABCB1^P1008G^ and ABCB1^E1007A^ lines was not different from *abcb1* mutants and *abcb1* complemented with Wt ABCB1, respectively (Fig. 1c-d). While as reported before (Hao et al. 2020), ABCB1^P1008G^ and ABCB1^E1007A^ were unable to restore IAA export defects of *abcb1* (Fig. 1g), ABCB1^P1008G^ and ABCB1^E1007A^ surprisingly revealed BL export not different from Wt ABCB1 (Fig. 1h). Importantly, export of the diffusion control BA (Suppl. Fig. 1b) and expression of mutant ABCB1 (Fig. 1; Suppl. Fig. 1h) was unchanged in comparison to Wt ABCB1, excluding indirect effects. To verify this apparent unilateral substrate discrimination of ABCB1^P1008G^, we expressed ABCB1^P1008G^ under its own promoter in the *abcb1 abcb19* double KO showing a prominent dwarfed shoot phenotype (Geisler et al. 2005; Noh et al. 2001; Effendi et al. 2015; Blakeslee et al. 2007; Bouchard et al. 2006). Surprisingly, all five independent *ABCB1:ABCB1^P1008G^* lines in the *abcb1 abcb19* background showed a nearly complete rescue of the shoot phenotype (Fig. 1i), which was not different from that the Wt ABCB1 version as exemplified by quantification of rosette leaf areas (Fig. 1j). These data argue for the fact that the *abcb1 abcb19* dwarfism is predominantly caused by BR transport defects, while auxin seems to have only a minor contribution, as shown here for the slightly reduced onset of inflorescences described for *abcb1* and *abcb19* single KOs (Geisler et al. 2005).

In summary, this dataset verifies a dual IAA and BL substrate specificity, which is limited obviously only to ABCB1 and ABCB19, and indicates that E-P1008 has an impact on the IAA but not BL transport capacity of ABCB1.

To uncover the molecular reason for this obvious unilateral substrate discrimination of ABCB1^P1008G^, we obtained a high-resolution cryo-EM structure of the ABCB1^P1008G^ mutant. To do so, ABCB1 was co-expressed with TWD1 in human embryonic kidney 293F (HEK293F) suspension cells to a large scale and purified for cryo-EM analysis in the presence of IAA ((Wei et al. 2025); Suppl. Fig. 2a-d). After data collection and processing, 3D reconstruction maps were obtained with an overall resolution of 3.8 Å (2e). Structural alignments with the Wt ABCB1 in its apo state (Ying et al. 2024) revealed an overall nearly identical structure (Fig. 2a-b) with only very subtle differences even for the E-P1008 loop (Fig. 2c).

**Fig. 2:**
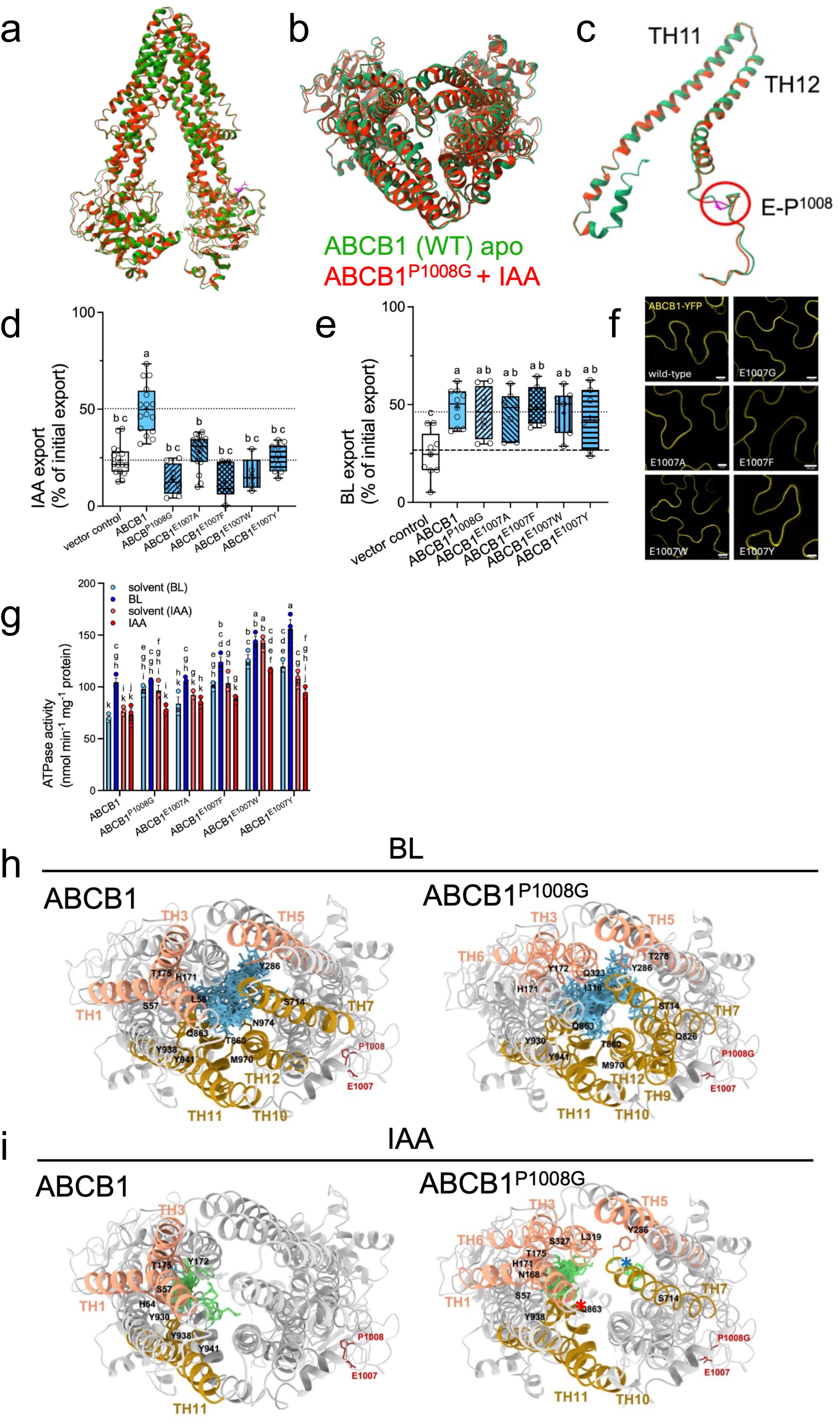
P1008A mutation of ABCB1 has no obvious effect on the overall ABCB1 structure but affects selectively IAA but not BL docking. **a-c**. Cryo-EM structural comparison of the Wt (apo) ABCB1 (green) and ABCB1^P1008G^ (red) in IAA-bound states. Cartoon representation of the entire ABCB1 structures (**a**), the NBD2 (**b**) and TH11 and TH12 as extension of E-P1008 (**c**). **d-f**. IAA (**d**) and BL (**e**) export from *N. benthamiana* protoplasts after transfection with indicated E-P1008 mutant versions of ABCB1. Significant differences (*p* < 0.05) of means ± SE (n ≥ 8 independent protoplast preparations) were determined using Ordinary One-way ANOVA (Tukey’s multiple comparison test) and are indicated by different lowercase letters. (**f**) Imaging of E-P1008 mutant versions of ABCB1 in in *N. benthamiana*; bar, 10 μm. **g.** ATPase activities of Wt and indicated E-P1008 mutant versions of purified ABCB1 in the absence (apo) and presence of indicated concentrations of BL and IAA. Data are means of three measurements ± SD. **h-i**. *In silico* docking (Autodock Vina) of BL (**i**) and IAA (**j**) to Wt ABCB1 (left) and ABCB1^P1008G^ cryo-EM structures (right). In a top view, helices from the Transmembrane Domain1 (TMD1) are coloured in pink and helices from TMD2 in brown; BL is in blue and IAA in green. A red asterisk marks absence of an IAA cluster near Y941 (TH11) and a blue asterisk marks the presence of a new IAA cluster between TH5 and TH7 in ABCB1^P1008G^.

Based on inward- and outward-facing structural models of ABCB1 (Hao et al. 2020) and inward-facing cryo-EM structures of ABCB1 (Wei et al. 2025; Chen et al. 2025), the E1007-P1008 peptide bond was found always in *trans*. We proposed thus that TWD1 might isomerize the E1007-P1008 peptide bond from *trans* to *cis* leading to enhanced IAA export (Hao et al. 2020; Geisler and Hegedus 2020). To sterically alter the E1007-P1008 equilibrium, we mutated E1007 in ABCB1 to aromatic amino acids thought to shift peptidyl-prolyl bonds to the *cis* conformation. Analysis of IAA export after heterologous expression in tobacco revealed that all three aromatic mutations completely blocked IAA export like found for ABCB1^P1008G^ or ABCB1^E1007Y^ (Fig. 2d). In contrast, all three aromatic mutations not significantly affected BL transport (Fig. 2e). As before, mutation of E1007 did not alter significantly transport of BA (Suppl. Fig. 1g).

Substrate-stimulated ATPase activity is an often used but unreliable metric for identifying true substrates because ABC transporters (including the ABCB subfamily) frequently exhibit uncoupled or poorly coupled mechanics (Livnat-Levanon et al. 2016). As shown before (Wei et al. 2025), the basal ATPase activity of ABCB1 remained unaffected in the presence of even 50 μM IAA but was stimulated by 5 μM BL (Fig. 2g). Subsequently, we quantified the effect of E1007-P1008 mutations on ATPase activities using purified ABCB1 protein (Fig. 2g; Suppl. Fig. 2f). Aromatic substitutions increased the basal ATPase activity suggesting that ATPase activity was uncoupled from IAA transport. Interestingly, ATPase activity of ABCB1^P1008G^ and ABCB1^E1007W^ was slightly inhibited by IAA. In agreement with intact BL transport, aromatic substitutions retained BL-stimulation suggesting a higher coupling efficiency for BL.

In order to exclude that absence of IAA-stimulation of ATPase activity was due to the absence of plant-specific factors, we quantified IAA and BL stimulation of vanadate-sensitive ATPase activity at on isolated microsomes prepared from transfected tobacco at pH 9, where ATPase activity of H^+^-ATPases is neglectable, excluding indirect effects (Aryal et al. 2023). Even though ABCB1 ATPase activity in the absence of substrate was only mildly higher than the vector control (Suppl. Fig. 1c), ATPase activities of ABCB1 microsomes were likewise stimulated by IAA than by BL at 5 μM, although the effect with IAA was not significantly different from the solvent control. Until now, it remains unclear whether IAA fails to significantly stimulate ATPase activity due to its lower affinity, smaller size or its dependence of an allosteric regulator.

Despite low-affinity binding of IAA to ABCB1 (Chen et al. 2025), IAA could not yet be traced in ABCB structures (Ying et al. 2024; Chen et al. 2025). Therefore, we investigated the potential of IAA and BL to bind to Wt and P1008G cryo-EM structures of ABCB1 by using *in silico* docking employing AutoDock Vina (Eberhardt et al. 2021). In Wt ABCB1, BL was bound as a single, central cluster filling nearly entirely the central substrate cavity formed by TH1, TH3, TH5, TH7 and TH10-12 (Fig. 2h; Suppl. Fig. 2e). This binding pose agrees with previous cryo-EM structures and identified residues (like Y286, Y941 and M970) that were identified and verified to be essential for BL transport via TH5, TH11 and TH12, respectively (Wei et al. 2025; Chen et al. 2025). In agreement with transport data, BL binding was widely conserved between Wt ABCB1 and ABCB1^P1008G^, except for a few contact residues for BL (like Q323 and N974) that were apparently lost (Fig. 2h). IAA docked to Wt ABCB1 as two main clusters with contacts provided by TH1,3 and TH11 (Fig. 2i; Suppl. Fig. 2e). Identified contact sites are in agreement with previously published IAA docking results using a modelled Arabidopsis ABCB1 structures (Bailly et al. 2011) but overlap also with identified BL-binding sites of Arabidopsis ABCB1 (Wei et al. 2025; Chen et al. 2025). Interestingly, IAA employed partially identical contact residues (like Y175, S57, Y938 and Y941) but also distinct ones (like Y172, H54 and Y930) provided by TH1,3,11. Importantly, the IAA cluster in proximity (ca. 8 Å) to Y941 of TH11 was entirely absent in ABCB1^P1008^ (Fig. 2h). Instead, a new IAA cluster was found where IAA is complexed by residues of TH5 and TH7 (like Y286), which is also involved in BL binding.

Taken together, these transport, ATPase and *in silico* data suggest that the conserved E-P1008 has a long distance-impact on ABCB1 activity and substrate recognition. They provide a structural and mechanistic ratio for the substrate discrimination of IAA but not of BL in the P1008G mutant of ABCB1, which is most likely caused by reduced IAA recognition and binding to the central substrate cavity.

To get a deeper insight into the mechanism of unilateral substrate discrimination by ABCB1^P1008^, molecular dynamics (MD) simulations of ABCB1 in solution at the coarse-grain (CG-MD) level of resolution were employed. A copy of the ligand (IAA, indole or tryptophan) under investigation was randomly placed in solution^1^ [^1^Unfortunately, we were unable to conduct CG-MD simulations with BL due to its molecular complexity.]. In 2 out of 5 replicates, IAA was binding to ABCB1 protein in its potential substrate binding site (or nearby; Fig. 3a). The binding occurred only when the protein was in its open conformation; simulations of the closed conformation using IAA as a ligand resulted in no binding (0 binding instances out of 5 replicas). Simulations of both the open and closed conformation of ABCB1 in presence of negative controls, either indole or tryptophan, resulted in no binding in 5 replicas, respectively (Fig. 3a), corroborating the experimental evidence (Bailly et al. 2008; Blakeslee et al. 2007; Geisler et al. 2005). Excitingly, likewise ABCB1^P1008G^ did not bind indole and tryptophan but also not IAA (0 binding instances out of 5 replica), which agrees with wet lab transport experiments (Fig. 1).

**Fig. 3:**
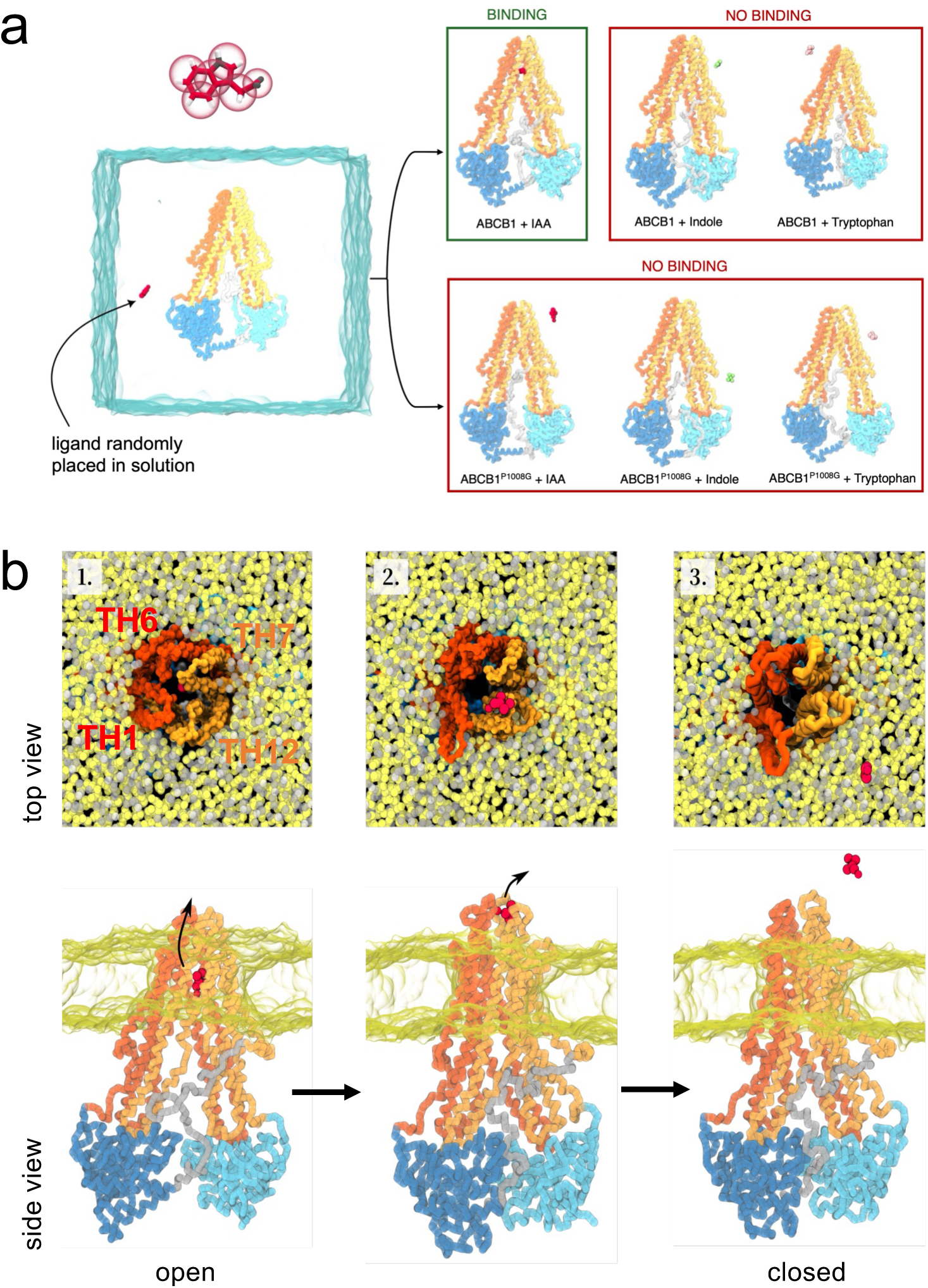
Coarse-grained molecular dynamics (CG-MD) provide a mechanistical explanation for IAA discrimination of ABCBP^1008G^. **a**. System setup of CG-MD simulation of ABCB1 in solution in presence of its substrate IAA (magenta); Inset, CG-MARTINI3 model of the IAA ligand (left). Representative snapshots from CG-MD simulations of ABCB1, both WT and P1008G mutant, placed in solution in presence of IAA (magenta), indole (green) and tryptophan (pink). MD simulations analyses reveal that IAA is the only substrate to successfully bind the presumed binding site in the WT-ABCB1 in its open conformation, while the same substrate fails to bind the E1007A-P1008G mutant. Both the negative controls indole and tryptophan did bind to the ABCB1 transporter. NBD1 and NBD2 are in dark and light blue, while TMD1 and TMD2 are in dark and light orange. **b.** Top (upper panels) and side views (lower panels) of CG-MD simulations representative frames overtime showing ABCB1 transporter embedded in a model membrane. The conformational change from open to close state allows the release of the substrate IAA (magenta) to the extracellular side.

CG-MD simulations were also employed to investigate the mechanism of opening and closing of ABCB1 employing the Switching-Go MARTINI model (Yang and Song 2024). Starting from a structure extracted from the unbiased MD simulation of open ABCB1 in solution described above, ABCB1 bound with IAA in its binding site was embedded into a pure POPC model membrane, and then the system was placed in water. The simulations consisted of 3 parts: i. a short simulation of the open ABCB1 in membrane with auxin bound; ii. a short MD relaxation to enable the switching to the closed conformation and iii. MD production of the closed conformation. In the first 2 steps, the IAA was retained into the ABCB1 structure, keeping its initial position in its binding site (Fig. 3b; Suppl Fig. 3; Suppl. Movie 1). During the 3rd step, when the protein conformation switched from open to closed, IAA was released into the extracellular space after only a few nanoseconds. Interestingly, a slight opening of the 4 helices, TH1, TH6, TH7 and TH12, rapping the ligand was sufficient for its release, which was happening before ABCB1 had yet completed its closing. In summary, CG-MD simulations underline the high degree of substrate specificity of ABCB1 (Geisler et al. 2005) and support the substrate discrimination of IAA by ABCB1^P1008G^.

To demonstrate the apparent regulation of substrate specificity on ABCB1 by means of a putative PPIase activity provided by TWD1, we deleted the *bona fide* PPIase activity of TWD1 by introducing two independent point mutations in the FKBD of TWD1. By analogy to mammalian FKBPs (Fanghanel and Fischer 2004), Y74 and Y132 (corresponding to Y26 and Y87 of HsFKBP12 ((DeCenzo et al. 1996)) were mutated to alanine that were shown before to be essential for immunosuppressant drug binding and PPIase activity (Suppl. Fig. 4a; (DeCenzo et al. 1996; Fanghanel and Fischer 2004)). Co-expression with TWD1^Y74A^ and TWD1^Y132A^, respectively, did not activate IAA export by ABCB1 (Fig. 4a) but resulted in export at vector control level, which might be caused by the fact that PPIase-defect TWD1 was eventually competing for the tobacco-endogenous ortholog of TWD1. As expected, mutation of TWD1 had no major effect on ABCB1-mediated BL export (Fig. 4b). Also, mutation of Y74 and Y132 in TWD1 did not significantly alter ABCB1 expression (Fig. 4c), ABCB1-TWD1 colocalization (Fig. 4d) or ABCB1-TWD1 interaction, as elaborated by ABCB1-TWD1 BRET and FRET analyses in tobacco (Fig. 4e-f).

**Fig. 4:**
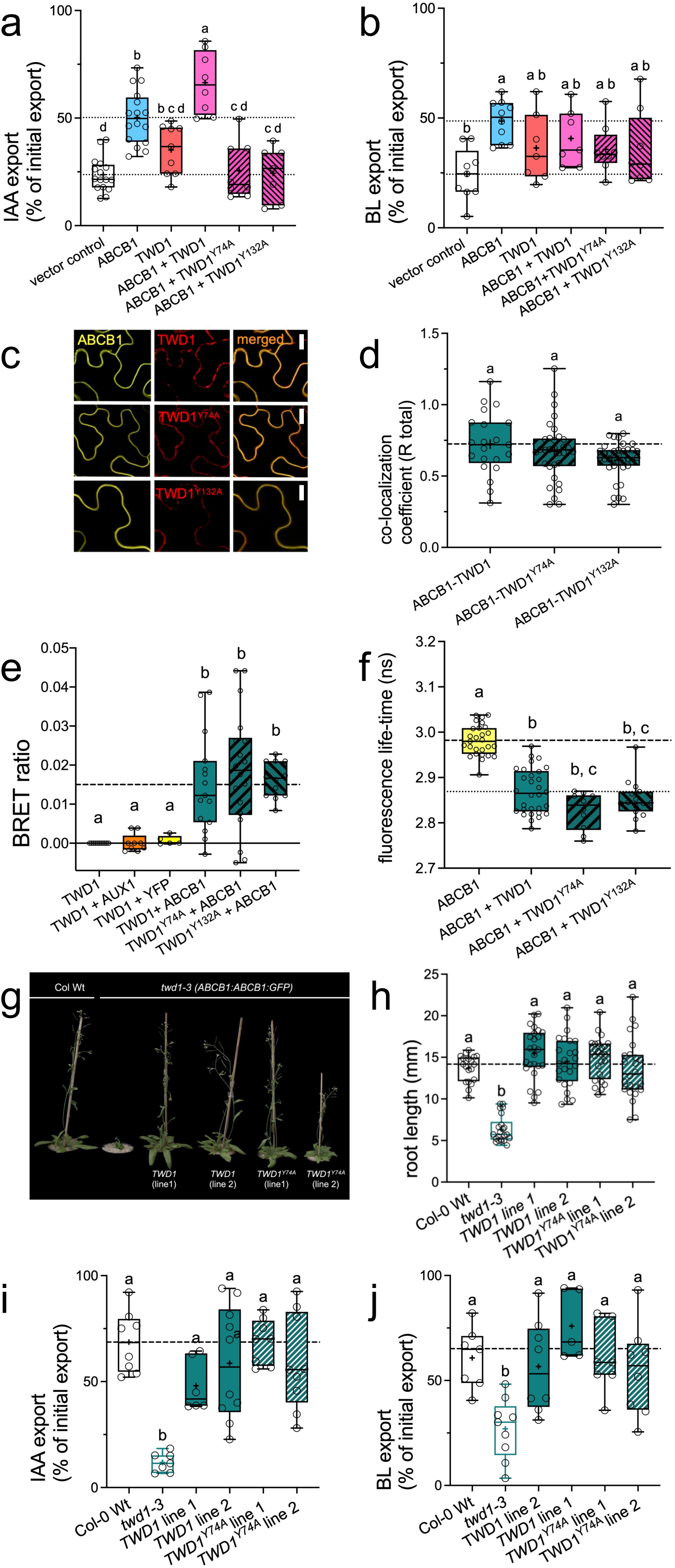
Activation of ABCB1-mediated IAA but not BL transport by TWD1 is dependent on a *bona fide* PPIase activity of TWD1. **a-b**. IAA (**a**) and BL (**b)** export from *N. benthamiana* protoplasts after co-transfection of ABCB1 with Wt and PPIase mutant versions (TWD1^Y74A^ and TWD1^Y132A^) of TWD1. Significant differences (*p* < 0.05) of means ± SE (n ≥ 8 independent protoplast preparations) were determined using Ordinary One-way ANOVA (Tukey’s multiple comparison test) and are indicated by different lowercase letters. **c-d.** Imaging (**c**) and co-localization of ABCB1 co-transfected with Wt TWD1, TWD1^Y74A^ or TWD1^Y132A^ tagged with YFP and mCherry, respectively, in tobacco (**d**); bar, 100 μm. **e-f.** ABCB1-TWD1 interaction analyzed by BRET (**e**) and FRET (**f**) using microsomes or leaf material from co-transfected tobacco, respectively. Significant differences (*p* < 0.05) of means ± SE (n ≥ 3 independent experiments) were determined using Ordinary One-way ANOVA (Tukey’s multiple comparison test) and are indicated by different lowercase letters. **g-h.** A PPIase-deficient (Y74A) version of TWD1 does complement growth defects of *twd1*. Phenotype of 30 dag pot-grown plants (**g**) and root length of 7 dag seedlings (**h**); bar, 2 cm. **i-j.** IAA (**i**) and BL (**j**) export from Arabidopsis *twd1-3* protoplasts complemented with Wt (TWD1) and PPIase-dead (TWD1^Y74A^) versions of TWD1. Significant differences (*p* < 0.05) of means (indicated by “+”) ± SE (n ≥ 6 independent protoplast preparations) were determined using Ordinary One-way ANOVA (Tukey’s multiple comparison test) and are indicated by different lowercase letters.

Next, we genetically complemented the *twd1-3* mutant allele of *TWD1* expressing *ABCB1-GFP* under its native promoter (Wu et al. 2010) with *TWD1^Y74A^.* PPIase-deficient TWD1 was widely able to complement the pleiotropic “*twisted dwarf1* syndrome” as indicated by restored overall plant stature (Fig. 4g), root (Fig. 4h; Suppl. Fig. 4d) and hypocotyl lengths (Suppl. Fig. 4c, e) and root gravitropism (Suppl. Fig. 4b). While TWD1^Y74A^ was able to complement shorter roots and hypocotyls of *twd1* to Wt or *twd1* complemented with Wt TWD1 level under standard conditions, this was interestingly not the case when plants were grown on high (1 μM) concentrations of IAA or BL (Suppl. Fig. 4d-e). In agreement with an overall complementation of the *twd1* mutant, TWD1^Y74A^ complemented also IAA and BL transport defects of *twd1-3* to Wt and Wt *TWD1* (*twd1*) levels (Fig. 4i-j). Complementation of growth and transport suggests that auxin and BR transporting ABCBs whose biogenesis is under control of TWD1 (Tsering et al. 2024) can reach the PM in *TWD1^Y74A^*lines. This co-chaperone function of TWD1 was shown to be provided by the tetratricopeptide repeat (TPR) domain of TWD1 (Tsering et al. 2024; Geisler et al. 2003) and thus to be independent of its PPIase activity. This is supported by confocal imaging showing that ABCB1-GFP indeed reaches the PM in Wt *TWD1* and *TWD1^Y74A^* lines (Suppl. Figure 3f). Label-free, quantitative proteomics suggest that only ABCB1,4,19,21 are chaperoned by TWD1 and that expression of all four is restored likewise in Wt *TWD1* and *TWD1^Y74A^* lines (Suppl. Figure 4f-g). In summary, this dataset suggests that disruption of the PPIase activity of TWD1 prevents the ability of TWD1 to activate ABCB1-mediated IAA transport above basic level but restores the growth defect of *twd1* to Wt level by providing basic IAA and BR transport.

Mutagenesis of the EP motif of ABCB1 and the FKBD of TWD1 suggested that a PPIase activity provided by TWD1 is responsible for altering substrate specificity of ABCB1 by *cis-trans* isomerization of the E1007-P1008 bond. Considering reported calmodulin binding to TWD1 (Kamphausen et al. 2002) and the finding that PPIase activity of FKBP38, the human ortholog of TWD1/FKBP42, was masked in the absence of calmodulin (Edlich et al. 2007b), we tested the calmodulin-dependence of TWD1 activity as a chaperone. First, we quantified the TPR-dependent intrinsic chaperone activity of purified TWD1 that is thought to be independent of its PPIase activity (Kamphausen et al. 2002). In light of the recently described TWD1-HSP90 interaction (Tsering et al. 2024), we posed the question, whether there is functional interdependence of HSP90 and TWD1 chaperone activities, as shown for mammalian multi-domain immunophilins (Iki et al. 2012). Monitoring protection against thermal aggregation of citrate synthase (CS) (Kamphausen et al. 2002) in the presence of TWD1 and HSP90 either alone or combined, revealed a higher chaperone (holdase) activity of HSP90 compared to TWD1, while intermediate CS aggregation kinetics of the combination at an equal total chaperone concentration indicated no apparent cooperativity (Fig. 5a-b). Next, we repeated the assay in the presence of Ca^2+^/calmodulin and found that, while holdase activity was only mildly induced in case of TWD1 and unchanged in case of HSP90, the combination of TWD1 and HSP90 was as efficient as HSP90 assayed alone in preventing CS aggregation, suggesting synergistic induction of a cooperative chaperone activity.

**Fig. 5:**
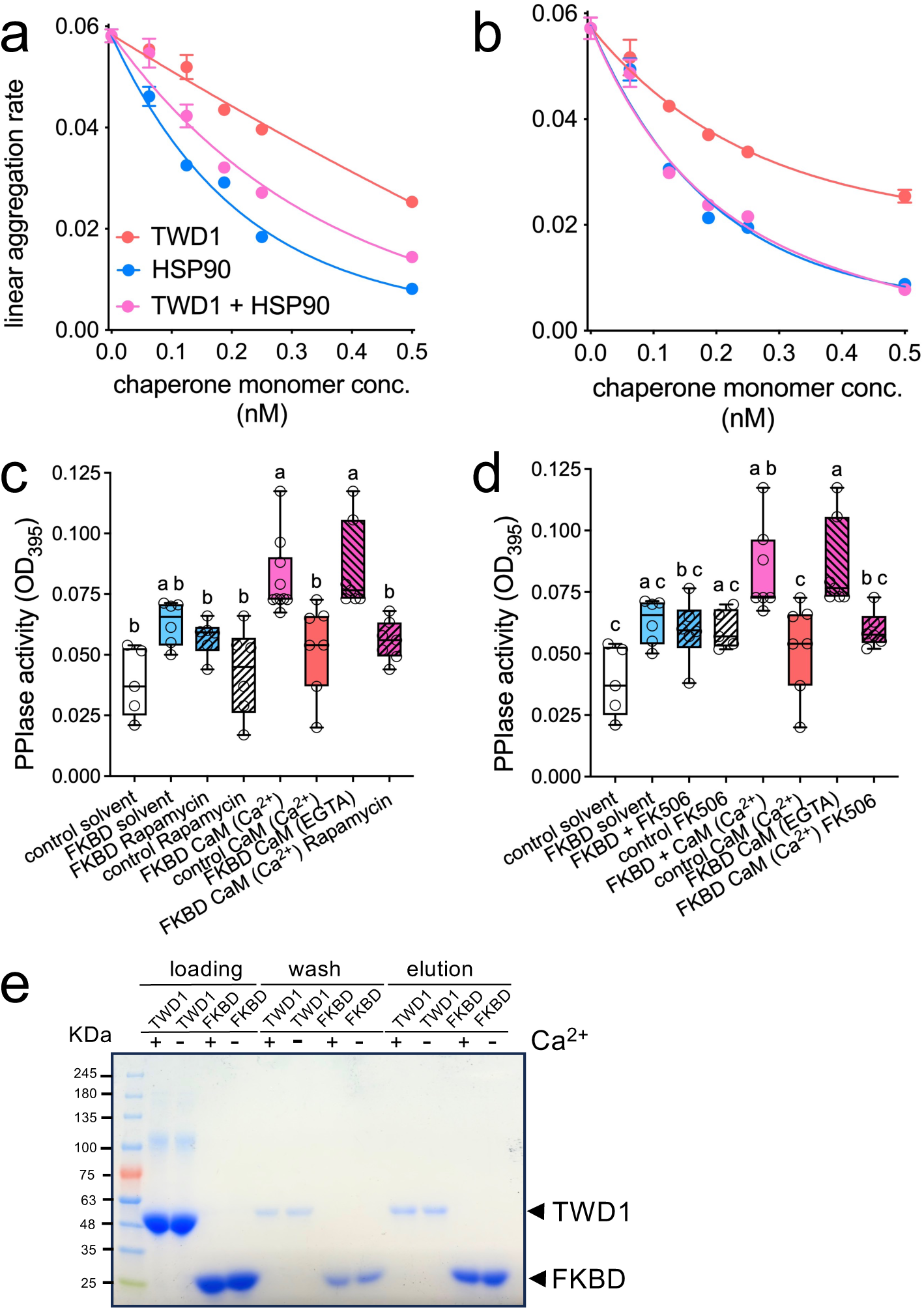
TWD1 owns a PPIase activity that is activated by calmodulin binding to the FKBD. **a-b**. Analysis of protection against temperature-induced citrate synthase aggregation (holdase) at 42°C in the presence of equimolar total chaperone concentration of TWD1 and bovine HSP90. Note that the shift of TWD1 + HSP90 in the presence of Ca^2+^/calmodulin suggests induced cooperativity; n=3-10. **c-d.** PPIase activity of the FKBD (TWD1^1-180^) was measured as protease-coupled standard PPIase in the presence and absence of the PPIase inhibitors FK506 (**c**) and rapamycin (**d**), respectively, as well as in the presence (1 mM Ca^2+^) or absence of calcium (5 mM EGTA). Significant differences (*p* < 0.05) of means (indicated by “+”) ± SE (n ≥ 8 independent PPIase measurements with two independent protein batches) to control (-TWD1 protein) were determined using Ordinary One-way ANOVA (Tukey’s multiple comparison test) and are indicated by different lowercase letters. **e.** The FKBD (aa 1-180) and full-length TWD1 minus membrane anchor (aa 1-339) bind to CaM-sepharose. Coomassie stain of equal volumes of FKBD and TWD1 protein taken form binding (total), washing and elution steps. Binding and washing steps were performed in the presence (1 mM Ca^2+^) or absence of calcium (5 mM EGTA).

Next, we measured the PPIase activity of recombinant FKBD (TWD1^1-180^) protein (Weiergraber et al. 2006) in the presence and absence of Ca^2+^/calmodulin using the standard PPIase tetrapeptide substrate, succinyl-AFPF-4-nitroanilide. PPIase activity of TWD1 without Ca^2+^/calmodulin was higher than the control but was not significantly inhibited by the immunosuppressant drugs and PPIase inhibitors, FK506 or rapamycin (Fig. 5c-d), a hallmark of FKBP PPIase activities (Fanghanel and Fischer 2004; Geisler et al. 2016; Geisler and Bailly 2007). In contrast, considerable rate acceleration of the TWD1 PPIase activity was observed in the presence of Ca^2+^/ calmodulin (Fig. 5c-d). Excess of FK506 or rapamycin competed with the substrate for binding to the PPIase site and thus decreased the rate of the *cis*-to-*trans* isomerization of the substrate down to the rate of spontaneous interconversion (Fig. 5c-d). Interestingly, TWD1 PPIase activation by calmodulin was not dependent on Ca^2+^ and thus also found in the presence of access of EGTA. Neither FK506/rapamycin nor Ca^2+^/calmodulin alone significantly promoted TWD1 activity, indicating that only the heterodimeric FKBD/calmodulin complex is the active PPIase form of TWD1. To demonstrate calmodulin binding to the FKBD, we performed binding assays of TWD1 (TWD1^1-339^) and FKBD (TWD1^1-180^) to calmodulin sepharose (Merck, Switzerland). Results indicate that FKBD and TWD1 protein both bind calmodulin (Fig. 5e). While FKBD binding seems independent of Ca^2+^, the TWD1 protein seems to be at least partially dependent on Ca^2+^ as washing and elution steps contain slightly more and less protein, respectively, in the presence of EGTA (Fig. 5e). Together, these data establish that intrinsic PPIase activity of TWD1 provided by the FKBD is functionally modulated by calmodulin, however, in an action independent of Ca^2+^.

To assess the mechanical effect of the P1008G mutation, we performed pulling simulations on the isolated intracellular loop containing residue P1008. During pulling, the force required to maintain the pulling velocity was recorded, and forces were extracted at defined end-to-end distances and averaged across simulations. At low extension, when the loops remained relatively relaxed, no significant difference was observed between the Wt and P1008G systems (Fig. 6a). However, once the distance between the two loop termini exceeded approximately 30 Å, the Wt loop required significantly higher pulling forces than the P1008G mutant. This indicates that the proline-containing Wt loop is mechanically more resistant to extension, whereas replacement by glycine increases local flexibility and reduces the force required to elongate the loop.

**Fig. 6:**
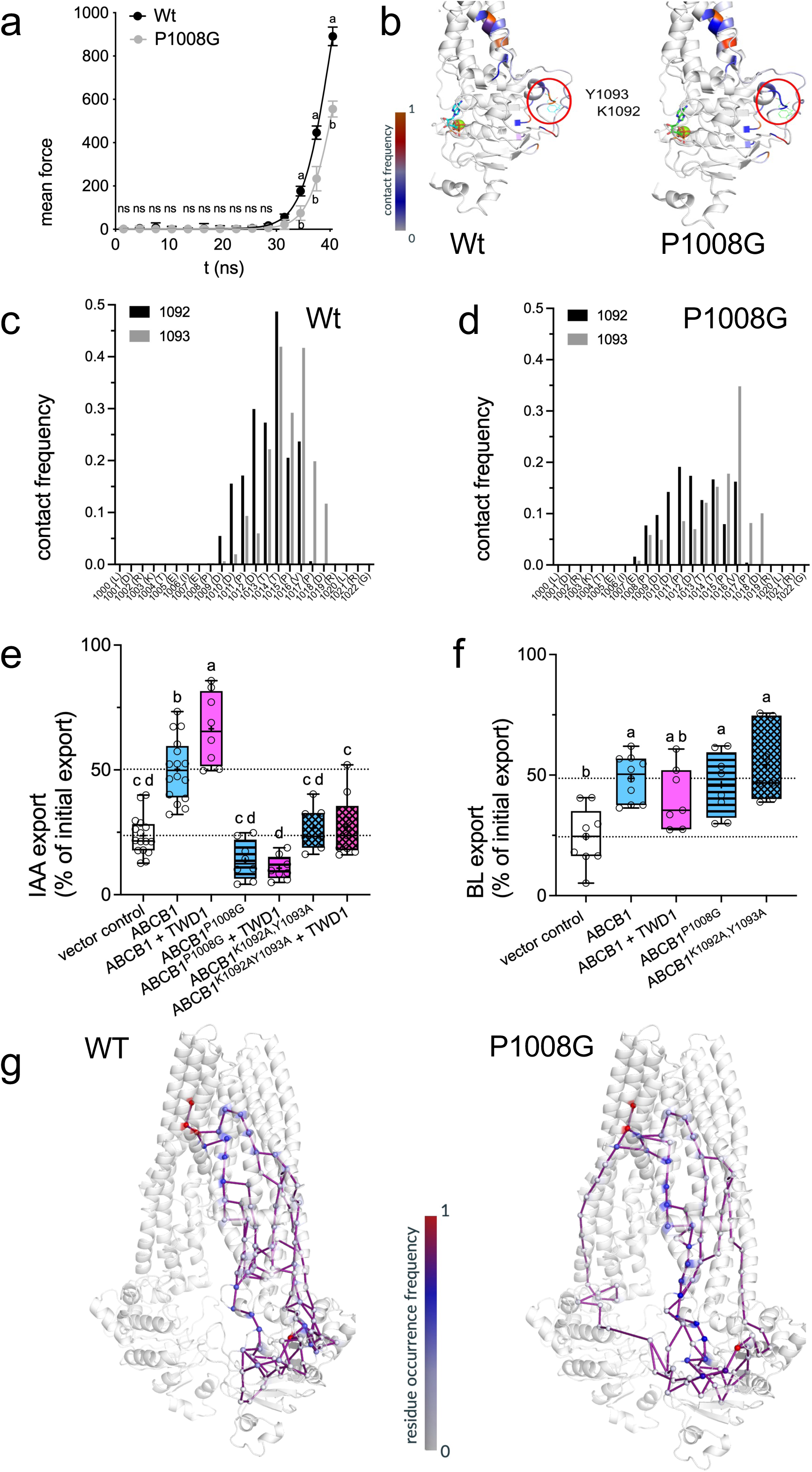
The P1008 loop communicates with the substrate-binding domain through intramolecular contacts involving K1092 and Y1093. **a**. Mechanical stiffness of the WT and P1008G ABCB1 loop region encompassing residues 1000-1022, assessed by incremental stretching simulations. The P1008G substitution reduces the stiffness of this loop region. **b.** Representative snapshots from full-length all-atom equilibrium MD simulations of Wt and P1008G ABCB1, highlighting the spatial proximity between the E1007/P1008 or E1007/G1008 loop region and the neighboring loop containing K1092 and Y1093 (red circles). Contact frequencies are mapped onto the structure. **c–d.** Quantification of residue-level contact frequencies between K1092 or Y1093 and the E1007/P1008-region in Wt ABCB1 (**c**) and the corresponding E1007/G1008-region in ABCB1-P1008G (**d**), as identified from full-length equilibrium MD simulations. **e–f.** IAA (**e**) and BL (**f**) export from *N. benthamiana* protoplasts after transfection with ABCB1 variants carrying K1092A and/or Y1093A substitutions. Significant differences (*p* < 0.05) among means ± SE (*n* ≥ 8 independent protoplast preparations) were determined by ordinary one-way ANOVA followed by Tukey’s multiple-comparison test and are indicated by different lowercase letters. **g.** Allosteric communication pathways between the NBD surface residue Y1093 and the substrate-binding pocket residue Y941 in Wt and P1008G ABCB1, derived from full-length all-atom equilibrium MD simulations. Spheres indicate residues along the inferred pathways; color coding shows residue occurrence frequency across the pathway ensemble. The P1008G mutation alters the preferred communication routes and reduces pathway redundancy.

Equilibrium simulations of the full-length Wt and P1008G ABCB1 proteins further supported a mutation-dependent rearrangement of local interactions. We first identified contacts between the P1008-containing loop and the rest of the protein. This analysis revealed reduced interactions of the loop with the NBD surface residues K1092 and Y1093 upon P1008G mutation (Fig. 6b). We then specifically analyzed the contacts formed by K1092 and Y1093 with residues of the P1008 loop. In the WT protein, these residues showed the highest contact frequencies with T1014 and V1016, respectively (Fig. 6c). In contrast, the P1008G mutant displayed weaker and more redistributed interactions, although the V1016-Y1093 contact remained the dominant interaction (Fig. 6d). These results suggest that the P1008G mutation alters the way in which the loop conformation is communicated to the NBD surface, potentially influencing conformational coupling toward the substrate-binding pocket. As a proof of concept, we quantified IAA and BL transport for the K1092A/Y1093A mutant versions of ABCB1. Like ABCB1^P1008G^, also ABCB1^K1092A,Y1093A^ was unable to transport IAA, while BL export was not affected (Fig. 6e-f). Also, TWD1 could not promote IAA transport of ABCB1^P1008G^ and ABCB1^K1092A,Y1093A^ (Fig. 6e-f).

To assess whether these local changes affect long-range allosteric communication, we analyzed potential allosteric pathways between the NBD surface residue Y1093 and the substrate-binding pocket residue Y941. The equilibrated parts of the trajectories were divided into overlapping 200-ns segments, and pairwise mutual-information-based motion correlations were calculated for each segment. These correlations were used to build weighted residue-interaction graphs, in which amino acids were represented as nodes and edges were weighted by their correlated motions. This pathway analysis suggested that the P1008G mutation alters the preferred communication routes between the NBD surface and the substrate-binding region. In particular, the mutant appeared to rely on a smaller number of dominant pathways (Fig. 6g). Such reduced pathway redundancy may make the communication between the loop, the NBD, and substrate-binding pocket more sensitive to regulatory perturbations. Together, these *in silico* findings suggest that the loop forms a mechanically and allosterically sensitive regulatory element, whose conformational dynamics can be tuned by calmodulin-activated TWD1 PPIase activity to modulate communication between the NBD surface and the substrate-binding pocket, providing substrate specificity alteration.

## Discussion

This experimentally demonstrated IAA transport activity of ABCB1 and ABCB19 (Chen et al. 2023; Ofori et al. 2018; Xu et al. 2014; Kubes et al. 2012; Kamimoto et al. 2012; Yang and Murphy 2009; Titapiwatanakun et al. 2009; Bailly et al. 2008; Bouchard et al. 2006; Geisler et al. 2005) has been recently challenged by four publications that reported BR transport for Arabidopsis ABCB1 and ABCB19 and cryo-EM structures in BL-bound states (Ying et al. 2024; Chen et al. 2025; Wei et al. 2025; Liu and Liao 2025). Moreover, three of the four studies failed to demonstrate IAA density in the cryo-EM maps (Ying et al. 2024; Chen et al. 2025; Wei et al. 2025).

Here, we provide conclusive biochemical evidence that ABCB1 harbors a dual auxin (IAA) and BR (BL) transport capacity (Fig. 1a-d). We report roughly five-times higher IAA transport rates for IAA than for BL and lower competition for IAA and BL to compete for IAA transport than for BL (Fig. 1c-d), respectively. These data agree with recent findings reporting lower IAA transport and binding affinities for IAA than for BL (Chen et al. 2025) employing, however, protein sources from mammalian cell lines obviously lacking plant specific factors. Interestingly, with ABCB1 and ABCB19 only two out of 11 ATAs from Arabidopsis seems to transport BL beside IAA (Fig. 1e-f). This is supported by complementation of the shoot dwarfism in *abcb1 abcb19* to nearly Wt employing an IAA transport-dead version of ABCB1 (Fig. 1g-j).

In summary, it appears that ABCB1 and ABCB19 are dual IAA/BR transporters that transport both substrates with different affinities and capacities, though with high degree of specificity (Blakeslee et al. 2007; Bouchard et al. 2006; Geisler et al. 2005). The latter is supported by MD analyses of ABCB1 transport (Fig. 3a). Currently it is unclear how BRs, as large, widely hydrophobic substrates, and IAA, as a small, hydrophilic ligand, binding both to widely overlapping clusters of the substrate binding pocket (Fig. 2 i-j; (Ying et al. 2024; Wei et al. 2025; Chen et al. 2025; Liu and Liao 2025)) are recognized. Interestingly, the BL/IAA binding pocket of ABCB1,19 is similar to the one of taxol in human ABCB (PDB6QEX; (Chen et al. 2025)), highlighting the conservation of substrate recognition between an ABCB functioning as MDR pump (Khunweeraphong and Kuchler 2021), and a highly specific, dual activity plant ABCB.

ATAs contain a signature D/E-P motif in a surface-exposed loop that was shown to be essential for IAA transport ((Hao et al. 2020); Fig. 1). The D/E-P is part of the previously mapped contact site between the FKBD of TWD1 and the NBD2 of ABCB1, respectively (Geisler et al. 2003). Here we provide multiple evidence that this motif is folded by a PPIase activity that is most likely provided by TWD1 to prioritize IAA over BL transport. First, mutation of both the acidic moiety as well as the adjacent proline disrupts IAA transport (Fig. 1g, Fig. 3a) but has no effect on BL transport (Fig. 1h). Second, TWD1 co-expression accelerates IAA but not BL transport (Fig. 4a-b). And third, disruption of the PPIase activity of TWD1 prevents TWD1’s ability to boost IAA transport (Fig. 4a-b). This proposed PPIase activity is provided by the FKBD of TWD1 (Fig. 5c-d), which in analogy to FKBP38 (Geisler and Hegedus 2020; Banasavadi-Siddegowda et al. 2011; Edlich et al. 2005) is activated by calmodulin binding to the FKBD itself (Kamphausen et al. 2002). Interestingly, also in analogy to FKBP38, calmodulin binding and activation of PPIase activity to the FKBD is independent of calcium (Fig. 5), while calmodulin binding to the TPR requires calcium (Kamphausen et al. 2002).

Substrate docking to the cryo-EM structures of ABCB1 using AutoDock suggests that unilateral IAA transport discrimination of IAA by ABCB1^P1008G^ is caused by loss and/or rearrangement of IAA but not of BL binding to the central substrate cavity of ABCB1^P1008G^ (Fig. 2h-i). A puzzling question was how this apparent substrate discrimination by *cis-trans* isomerization is mechanistically transferred from the D/E-P motif to the substrate binding site. The E-P motif of ABCB1 is in direct connection with TH12 reaching toward the substrate binding pocket (Fig. 2h-i). However, the subtle steric differences between Wt ABCB1 and ABCB1^P1008G^ proteins (Fig. 2), together with the fact that proline and glycine are both uncharged and that this motif lies within a flexible loop, argue against a direct steric or charge transfer to the substrate binding domain as mechanistic explanation. Full ABCB1 equilibrium MD simulations revealed reduced intramolecular contacts in ABCB1-P1008G between residues around E1007/G1008 and a spatially adjacent residues K1092 and Y1093. This effect is most likely caused by increased flexibility of the E1007–G1008 loop in the P1008G mutant (Fig. 6a-d). Reduced IAA but intact BL transport for the K1092A/Y1093A mutant versions of ABCB1 support this scenario (Fig. 6e-f).

An emerging aspect of our work is that dual substrate ABCBs, like ABCB1 and ABCB19, function as integrators of auxin and BR transport pathways (Geisler 2024). A “switch” between IAA/BR transport specificities is provided by the FKBP42, TWISTED DWARF1. Our *twd1* and *abcb1 abcb19* complementation data indicate that the PPIase activity of TWD1 (Fig. 4g-j) and IAA transport by ABCB1 (Fig. 1i-j), respectively, is not essential for normal plant growth during that ABCB1,19 would function primarily as BR transporters. However, in certain developmental or physiological programs requiring IAA transport, an activation of TWD1’s PPIase by calmodulin would occur. This activity would isomerize the D/E-P peptide bond, transferring ABCB1/19 into a high-affinity state for IAA, which prioritizes IAA over BR transport (Suppl. Figure 5). The connection between calcium, calmodulin and auxin transport has been demonstrated through extensive research, establishing a key regulatory role for calcium signaling in controlling the polarity of auxin efflux (Vanneste and Friml 2013). The beauty and novelty of such a mechanism of transient ABCB activation is that it would fall non-enzymatically back to its basal state after seconds to minutes (Fanghanel and Fischer 2004).

Importantly, this model provides also a molecular rational why all attempts to capture an IAA-bound state and demonstrate IAA-stimulation of ATPase activity for ABCB1/19 have failed so far. As reported here, in the absence of TWD1 ABCB1 owns indeed very low IAA rates (Fig. 1; (Chen et al. 2025; Geisler et al. 2005; Yang and Murphy 2009) and thus most likely also low IAA affinities. Although ABCB1/19 protein production requires co-expression with TWD1 (Ying et al. 2024; Chen et al. 2025; Wei et al. 2025; Liu and Liao 2025), purified ABCB1/19 can be expected to fall back to a low affinity state for IAA due to loss of TWD1. However, our data demonstrating that ATPase is activated in aromatic E1007 ABCB1 mutants and that IAA inhibits ATPase in aromatic E1007 ABCB1 mutants agree with the finding that IAA was shown to be able to cause a conformational change of ABCB1 into an outward-open conformation (Chen et al. 2025).

An exciting parallel between plant ABCB-TWD1 and human CFTR-FKBP38 modules is provided by the finding that FKBP38 has been implicated in CFTR/ABCC7 maturation and regulation (Geisler and Hegedus 2020; Banasavadi-Siddegowda et al. 2011; Edlich et al. 2005). However, so far, no essential proline-isomerization mechanism for CFTR channel opening has been definitively established. Interestingly, CFTR owns a structurally conserved and surface exposed proline (Geisler and Hegedus 2020), suggesting that such a regulatory mechanism might be evolutionary conserved. Moreover, Ca^2+^-bound calmodulin can directly bind and increase the CFTR open probability, providing complementary CFTR fine-tuning (Csanady et al. 2019).

## Acknowledgments

We would like to thank L. Charrier, M. Currat and M. Stumpe for excellent technical assistance. This work was supported by grants from the Swiss National Funds (project IZ11Z0_230926 and 320030-236105 to MMG) and the Hungarian National Research Development and Innovation Office Funds (K 137610, TKP2021-EGA-23, and HU-RIZONT-2024-00003 to TH).

## Author contribution

TT, FR, JS, MdD, AB, TH, NF, LS and MMG designed research; TT, FI, AB, MdD, JS, MdD, PH, HZ, XX and HW performed research; TT, FI, MdD, JS, AB, PR, HZ, XX, HW and MG analyzed data; SV, TH, JR, LD and MMG supervised work; MMG wrote the manuscript, all authors commented on the manuscript.

## Data availability

Requests for data should be made to and will be fulfilled by M.M. Geisler provided the data will be used within the scope of the originally provided informed consent. Source data are provided with this paper.

## Methods

### Plant Material and Phenotypic Analyses

The following *Arabidopsis thaliana* lines were used: *abcb1-1, abcb19-1*, *abcb1-1 abcb19-1*, *twd1-1* (Geisler et al. 2003); *ABCB1:ABCB1-GFP*, *ABCB1:ABCB1^P100G^-GFP* and *ABCB1:ABCB1^P100G^-GFP* (Hao et al. 2020). *35S:TWD1-mCherry and 35S:TWD1^Y74A^-mCherry* lines were created by site-directed mutagenesis using the QuikChange Lightning Site-Directed Mutagenesis Kit (Agilent) and transforming *twd1-3* (Geisler et al. 2003) by using floral dipping. Isogenic, homozygous lines for the transgene in the F3 generations were used for down-stream analyses.

Seedlings were generally grown on vertical plates containing 0.5 Murashige and Skoog media, 1% sucrose, and 0.75% phytoagar in the dark or at 8 h (short day), 16 h (long day), or 24 h (constant) light per day. Developmental parameters, such as root gravitropism and root or hypocotyl elongation, were quantified by imaging as described in (Wang et al., 2013). All experiments were performed at least in triplicate with 10 to 30 seedlings per each experiment.

### Protein-protein interaction analyses

FRET-FLIM analyses were performed as described elsewhere (Aryal et al. 2023). In short, indicated binary vectors and *p19* as gene-silencing suppressor were transformed into *Agrobacterium tumefaciens* strain GV3101 and infiltrated into *Nicotiana benthamiana* leaves. The measurements were performed 3 dai using a SP8 laser scanning microscope (Leica Microsystems) as described (Aryal et al. 2023).

For BRET analysis, *N. benthamiana* leaves were Agrobacterium co-infiltrated with indicated BRET construct combinations (or corresponding empty vector controls) and microsomal fractions were prepared 4days after inoculation (dai). BRET signals were recorded from microsomes (each ∼10 μg) in the presence of 5 µM coelenterazine (Biotium Corp.) using the Cytation 5 image reader (BioTek Instruments) and BRET ratios were calculated as described previously (Wang et al. 2013). The results are the average of 20 readings collected every 30 seconds, presented as average values from a minimum of three independent experiments (biological replica: independent Agrobacterium infiltrations and microsome preparations) each with four technical replicates.

### Auxin and brassinosteroid transport

^3^H-indol-3-acetic acid (^3^H-IAA; ARC ART0340, 25 Ci/mmol)), ^14^C-benzoic acid (^4^C-BA; ARC ART0186A, 55 Ci/mmol) and ^3^H-brassinolid (^3^H-BL; custom-synthesized by ARC, 56 Ci/mmol) export from *Arabidopsis* and tobacco mesophyll protoplasts was analyzed as described (Henrichs et al. 2012). Relative export from protoplasts was calculated from exported radioactivity into the supernatant as follows: (radioactivity in the supernatant at time t = x min.) - (radioactivity in the supernatant at time t = 0)) * (100%)/ (radioactivity in the supernatant at t = 0 min.); presented are mean values from >6 independent protoplast preparations.

Uptake of 1 nM ^3^H-IAA and ^3^H-BL into *Arabidopsis* vesicles prepared from Arabidopsis lines grown as mixotrophic liquid cultures was measured in the absence (solvent) or presence of 1000x access of non-labelled IAA or BL, respectively, as described in (Matern et al. 2019).

### Confocal laser scanning microscopy

For imaging, seedlings were generally grown for 5dag on vertical plates containing 0.5 Murashige and Skoog media, 1% sucrose, and 0.75% phytoagar at 16h (long day) light per day. For chemical treatments, seedlings were transferred for 12h on test plates containing the indicated chemicals or the solvents. For confocal laser scanning microscopy work, an SP5 confocal laser microscope was used. The following confocal settings were set to record the emission of GFP (excitation 488 nm, emission 500– 550 nm), YFP (excitation 514 nm, emission 524–550 nm) and mCherry (excitation 587 nm, emission 550–650nm).

### ABCB1 protein expression and cryo-EM analyses

HEK293F cells (Sino Biological) were cultured in SMM 293T-ll medium (M293TIl, Sino Biological) at 37°C and 130 rpm under 5%CO_2_ and transfected upon reaching a density of 2 x 106 cells per ml. A total of 1.5 mg of plasmids of ABCB1 and TWD1 at a ratio of 2:1 were pre-incubated with 4 mg of linear polyethylenimines (Polysciences) in 45 ml of medium for 15 min.The mixture was then added to 800 ml of HEK293F cells, followed by a static incubation for 15 min. After a 12 h transfection, 10 mM sodium butyrate (Sigma-Aldrich) was added to the cells , and they were cultured for an additional 48 h at 30°C. The transfected cells were harvested by centrifugation at 2100 *g* for 10 min, and the cell pellets were resuspended in lysis buffer containing 25 mM HEPES-KOH (pH 7.4) and 150 mM NaCl, to which protease inhibitor cocktail (1 mM PMSF,1.3 μg ml^-1^ aprotinin, 0.7 μg ml^-1^ pepstatin A, and 5 μg ml-1 leupeptin) and 1.5% (w/v) dodecyl maltopyranoside (DDM, Anatrace) supplemented with 0.3% (w/v) cholesteryl hemisuccinate (CHS, Sigma-Aldrich) were added. After incubation in a rotating shaker at 4°C for 2 h, the mixture was centrifuged at 200,000 *g* for 60 min. The supernatant was isolated and incubated with anti-FLAG M2 affinity resin(Sigma-Aldrich) at 4°C for 40 min(Wei et al. 2025).

For cryo-EM sample preparation, the resin was rinsed three times using lysis buffer plus 0.06%(w/v) digitonin (Apollo Scientific) and then eluted using lysis buffer supplemented with 0.06%(w/v) digitonin (Apollo Scientific) and 200 μg ml^-1^ FLAG peptide. The protein eluent was concentrated to 2 ml using a 100-kDa cutoff Centricon filter(Millipore) and further puri-fied by size-exclusion chromatography (Superose-6 Increase,10/300GL,GE Healthcare) in lysis buffer plus 0.03% (w/v) digitonin. Peak fractions were pooled together and further concentrated to 10 mg ml^-1^ before cryo-EM sample preparation. All purification procedures were performed at 4°C.

Cryo-EM sample preparation was performed in the presence of 10 mM IAA and incubated on ice for 30 min before cryo-EM sample preparation. Samples were prepared using the Vitrobot Mark lV (Thermo Fisher Scientific).In brief, a 4ul protein aliquot was applied to a holey carbon grid (Quantifoil Cu R1.2/1.3,300mesh) glow discharged by SOLARUS 950 Plasma Cleaner (Gatan) using H_2_ and O_2_ for 10 s. The grid was blotted with grade 597 Filter Paper(Cytiva Whatman) for 3 s at 8°C and 100% humidity. The grid was then plunged into liquid ethane precooled by liquid nitrogen and transferred to a storage box.

All cryo-EM data were collected with EPU software in super-resolution mode on a 300-kV Titan Krios microscope (Thermo Fisher Scientific) equipped with a K3 Summit direct electron detector(Gatan) and a GIF Quantum energy filter (Gatan) at a nominal magnification of 81 000× with defocus values ranging from -1.0 to -2.0 μm and a calibrated pixel size of 0.55 Å. Each movie stack was acquired with an exposure time of 3 s anddose-fractioned into 32 frames, yielding a total accumulated dose of 50 e^-Å-2^. Data were then imported into RELION 4.0, motion corrected, and dose weighted with MotionCor2 (Zheng et al. , 2017;Zivanovet al., 2018).The resulting micrographs were binned two-fold, yieldinga pixel size of 1.1 Å, and then imported into cryoSPARC (v.3.2.0) (Punjani et al., 2017). The defocus values of each image were determined using CTFFIND4 (Rohou and Grigorieff, 2015). A detailed procedure for ABCB1^P1008G^ is described below as an example.

For ABCB1^P1008G^, 3,204 micrograph stacks were collected. 3,167,663 particles were automatically picked by the template picker in cryoSPARC and then applied to multiple rounds of 2D classification. 2,887,401 particles were selected and subjected to *ab initio* reconstruction with six classes. C1 symmetry was applied during the entire process. 239,513 particles were selected for non-uniform refinement. An EM map at 4.1 Å was obtained with clear structural features and served as a reference for further heterogeneous refinement. After several rounds of heterogeneous refinement and *ab initio* reconstruction, 117,377 particles were selected for non-uniform refinement, finally yielding a reconstruction map at 3.8 Å resolution. The overall resolution of the final maps of ABCB1^P1008G^ were determined by the gold-standard Fourier shell correlation at a 0.143 criterion (Chenet al.,2013), and the local resolutions were estimated using implements in cryoSPARC.

The apo-state structure model of ABCB1 predicted by AlphaFold2 was docked into the obtained cryo-EM density maps using UCSF Chimera (Pettersen et al., 2004;Jumper et al., 2021). The fitted structures were manually adjusted in Coot (Emsley and Cowtan, 2004).Further refinements were performed using phenix. real_space_refine in PHENIX (Adams et al., 2010). The IAA molecule was built and refined using restraints generated by Elbow (Adams et al., 2010). All structural figures were prepared with PyMol or UCSF Chimera software.

### Coarse-grained MD simulations

MD simulations of the ABCB1 were performed at the coarse-grain (CG) level of resolution utilizing the cryo-EM structures of the WT and P1008G ABCB1, in both its open and closed conformations. Missing loops in the experimental structures were resolved using AlphaFold2 (AF3; (Jumper et al. 2021). The MARTINI3 CG-model (Souza et al. 2021) was employed along with the GROMACS software. The parameters for the auxin molecule were adapted from both the indole and tryptophan molecules.

To assess the spontaneous binding of the auxin ligand versus control molecules such as indole and tryptophan in both WT and E1007A-P1008G mutant ABCB1 structures, MD simulations of the protein in solution were performed, following a well-established protocol (Alvarez et al. 2025). Initially, the energy-minimized atomistic protein was mapped onto the MARTINI3 Elastic Network (EN) model using the martinize2 tool (Souza et al. 2021).The protein was then inserted, together with a single copy of the ligand (auxin, indole, or tryptophan), into a MARTINI water box containing 0.12 M NaCl. The system was energy-minimized using the steepest descent algorithm, followed by a short 60 ns isobaric–isothermal (NPT) equilibration using the Berendsen barostat (Parrinello and Rahman 1981)and the V-rescale thermostat (Bussi et al. 2007)

Production simulations were performed as five independent replicas for each system (WT/mutant ABCB1 + auxin, WT/mutant ABCB1 + indole, and WT/mutant ABCB1 + tryptophan), with each replica run for 2 μs, corresponding to a total simulation time of 60 μs overall. During production, the temperature was maintained at 310 K with a coupling time of 1 ps using the V-rescale thermostat, and the pressure was kept at 1 bar with a coupling time of 12 ps using the Parrinello–Rahman barostat (Parrinello and Rahman 1981).

To investigate the opening/closing mechanism of the ABCB1 protein while bound to a molecule of auxin, CG-MD simulations employing the Switching-Go MARTINI model (Yang and Song 2009) were performed. A representative snapshot of the Wt ABCB1 protein in its open conformation with a copy of the auxin molecule bound in its potential binding site was taken from the unbiased CG-MD simulations in solution described above. The complex protein-ligand was the embedded into a 1-palmitoyl-2-oleoyl-sn-glycero-3-phosphocholine (POPC) model membrane utilizing the *insane.py* script (Wassenaar et al. 2015), then water and 0.15 M NaCl were added. Following energy minimization using the steepest descent algorithm, the system was equilibrated in the NPT ensemble with semi-isotropic pressure coupling, using the V-rescale thermostat and Berendsen barostat to maintain temperature and pressure. Three successive MD simulation steps were then performed. First, a short MD simulation (*d*t = 20 fs) of the open conformation was carried out using a contact map derived from the open structure. Second, a short relaxation MD simulation (*d*t = 2 fs) was performed to promote the transition from the open to the closed conformation, using a contact map based on the closed structure. Third, a production MD simulation of the closed conformation (*d*t = 20 fs) was conducted using the corresponding closed-state contact map. Three replicas of 550 ns each were simulated. All contact maps were generated with the martinize2 tool using the Go-model framework.

Renders of the snapshoots deriving from MD simulations were generated using the Visual Molecular Dynamics (VMD) software (Humphrey et al. 1996).

### All-atom MD simulations

To generate inputs for the simulations, the apo ABCB1 structure was derived from the refined structural model ABCB1-J670_apo_3p5_final.pdb. For full-length simulations, the Wt ABCB1 structure was submitted to CHARMM-GUI/Membrane Builder for system assembly. The P1008G mutant was generated during the CHARMM-GUI membrane-protein preparation workflow [25130509]. Unless otherwise stated, protonation states, terminal patches, membrane embedding, solvation, ion placement and the initial equilibration protocol were assigned. CHARMM36m all-atom force field was used, and the system was solvated with explicit TIP3P water (Huang et al. 2017; Boonstra et al. 2016). The protein was embedded in a mixed lipid bilayer using the membrane composition applied throughout the study: POPC at a 1:1 ratio in the extracellular leaflet and POPC:PLPC:POPS at a 37:43:12:8 ratio in the intracellular leaflet. The systems were neutralized and supplemented with 150 mM KCl. Long-range electrostatics were treated with the particle-mesh Ewald method, and short-range non-bonded interactions were truncated at 1.2 nm. Bonds involving hydrogen atoms were constrained with the LINCS algorithm (Hess et al. 1997). Energy minimization and stepwise equilibration were performed using the CHARMM-GUI-generated GROMACS input files. Production simulations were performed with GROMACS 2025 under constant particle number, pressure and temperature conditions (Abraham et al. 2015). The temperature was maintained at 303.15 K, and pressure was maintained at 1 bar using semi-isotropic pressure coupling. A 2-fs integration time step was used for full-length equilibrium simulations. Equilibrium molecular dynamics simulations were performed for the full-length wild-type and P1008G ABCB1 systems. Each system was simulated in two independent 2 μs equilibrium runs initiated with different random velocities, resulting in a total simulation time of 4 μs per system. The first part of each trajectory was treated as relaxation, and the equilibrated 1000–2000 ns interval was used for contact and allosteric-pathway analyses. Before analysis with GROMACS tools and MDAnalysis (Michaud-Agrawal et al. 2011), trajectories were fitted to the protein backbone.

To assess the mechanical effect of the P1008G mutation independently of the full ABCB1 structure, biased pulling simulations were performed. The isolated P1008-containing loop was extracted from the wild-type and P1008G ABCB1 models. The loop peptide was oriented along its principal axis and placed into an elongated rectangular simulation box (box length of 18 nm along this axis) to accommodate extension during pulling. The peptide was solvated with TIP3P water and neutralize2d with KCl (150 mM) using standard GROMACS protocols implemented locally with GROMACS 2025 and the CHARMM36m. Biased production simulations were performed with GROMACS 2022.3 patched with PLUMED 2.8.1 (consortium 2019). Simulations were carried out in the NPT ensemble at 303.15 K and 1 bar using a velocity-rescaling thermostat and a C-rescale barostat (Bussi et al. 2007), LINCS for hydrogen bond constraining, and PME for long-range electrostatics. Loop extension was monitored using the end-to-end distance between the two loop termini as the collective variable. Constrained stretching simulations were performed by applying a moving harmonic restraint to this end-to-end distance with a force constant of 2000 kJ mol⁻¹ nm⁻². Each pulling simulation was run for 4.05 × 10⁷ steps, corresponding to 40.5 ns. Collective-variable and bias-force information was written every 5,000 steps. For comparative analysis, the pulling force was extracted from the PLUMED output and binned according to the instantaneous end-to-end distance. Forces at defined distances were averaged across simulations for the Wt and P1008G loop systems.

Contact maps were calculated from the equilibrated parts of the wild-type and P1008G trajectories using MDAnalysis in custom Python scripts. Contacts were defined between Cα atoms with a distance cutoff of 7.5 Å. Contact frequencies were calculated as the fraction of analyzed frames in which a given residue pair satisfied this distance criterion.

Allosteric pathway analysis was performed between Y1093 and Y941. The equilibrated 1000–2000 ns interval of each trajectory was divided into nine overlapping 200-ns segments: 1000–1200, 1100–1300, 1200–1400, 1300–1500, 1400–1600, 1500–1700, 1600–1800, 1700–1900 and 1800–2000 ns. This resulted in 18 segment-specific analyses in total for each construct. For each segment, pairwise residue motion correlations were calculated using the generalized correlation measure based on mutual information as implemented in Wordom (Seeber et al. 2011). Amino acids were represented as nodes in residue-interaction networks. Edges were assigned according to residue contacts and weighted by the mutual-information-based correlation coefficient. Edge weights were calculated as −log(Cij), where Cij is the generalized correlation between residues i and j. In this representation, highly correlated residue pairs have shorter effective graph distances. Shortest communication paths between Y1093 and Y941 were calculated for each segment-specific network. Pathway usage was then compared between Wt and ABCB1^P1008G^ by counting the residues and edges recurring across the segment-wise shortest paths.

Plots were generated with Matplotlib (Hunter 2021). Molecular structures were visualized using PyMOL (The PyMOL Molecular Graphics System, Version 2.5 Schrödinger, LLC).

### Measurement of PPIase and CS aggregation activity

PPIase activity of purified TWD1 (TWD1^1-339^) and TWD1 FKBD (TWD1^1-180^ ; (Scheidt et al. 2006)) was determined using a protease-coupled assay and succinyl-ALPF-4-nitroanilide as a substrate (Fanghanel and Fischer 2004) PPIase activity was measured in a reaction mixture containing 1 μM TWD1 protein, 5 μM recombinant bovine CaM and 2 mM CaCl_2_ or EGTA. In some cases, 5 μM FK506 (Fujisawa) or rapamycin (Merck) were added.

### Calmodulin binding assays

Binding of purified TWD1 (TWD1^1-339)^ and FKBD (TWD1^1-180)^ protein REF to Calmodulin-Sepharose^™^ 4B (Merck, Switzerland) was executed in columns according to the manufacturer. Binding and washing were performed in the presence (1 mM Ca^2+^) and absence of calcium (5 mM EGTA), respectively; elution was done by addition of prewarmed (95°C) 1x loading buffer. Equal volumes were separated by 4-20% PAGE (Biorad, Switzerland) followed by Coomassie blue staining.

### ATPase assay

ATPase activities of purified WT Arabidopsis ABCB1 proteins and its mutants were measured by using the ATPase Colorimetric Assay Kit (Innova Biosciences) as described recently (Ying et al. 2024).

ATPase activities from microsomes (0.06 mg/ml) prepared from tobacco plants transfected with vector control or WT ABCB1 using the colorimetric determination of ortho-phosphate released from ATP (Chifflet et al. 1988). Briefly, microsomes were added to ATPase buffer (20 mm MOPs pH 7.7 or pH 9.0, 8 mm MgSO_4_, 50 mm KNO_3_, 5 mm NaN_3_, 0.25 mm Na_2_MoO_4_, 2.5 mm phosphoenolpyruvate, 0.1% pyruvate kinase) in the presence and absence of 0.5 mM sodium ortho-vanadate. The reaction was started by the addition of 15 mm ATP and incubated at 37°C for 15 min with shaking. The amount of Pi released in the absence (solvent control) or presence of IBA, IAA, CLX, indole, or *ortho*-vanadate (50 μM) was quantified using a Cytation 5 reader (BioTek Instruments).

### In silico docking

IAA and BL were docked to the central cavity of ABCB1 using AutoDock Vina (Eberhardt et al. 2021) as part of the SwissDock platform (swissdock.ch) employing default settings. Results were analyzed in UCFS ChimeraX Vers. 1.8 (https://www.rbvi.ucsf.edu/chimerax).

### Statistics & Reproducibility

Data were statistically analyzed using Prism 11.0.1. (GraphPad Software, San Diego, CA). Normal (Gaussian) distribution of values was tested prior to statistical analyses using the Prism-embedded D’Agostino-Pearson omnibus normality test. For statistics, either “Ordinary One-way ANOVA“ or “Two-way ANOVA” and indicated comparison tests were used as post-hoc analysis to determine significances. Data are presented as “box-and-whisker plots”, where median and 25^th^ and 75^th^ percentiles are represented by the box itself and the middle line; means are indicated by a “+”.

No statistical method was used to predetermine sample size. No data were excluded from the analyses; the experiments were not randomized; the investigators were not blinded to allocation during experiments and outcome assessment.

## Supplemental Data Figures and Movies

**Supplemental Fig. 1: Transport and *abcb1* complementation controls**

**Supplemental Fig. 2: ABCB1 expression and docking controls**

**Supplemental Fig. 3: CG-MD simulation controls**

**Supplemental Fig. 4: *twd1* complementation controls**

**Supplemental Fig. 5: Speculative model on unilateral IAA transport activation of ABCB1 by TWD1**

**Suppl. Movie 1: Movie of CG-MD simulations showing ABCB1 transporter in solution.** The conformational change from open to close state of ABCB1 (side view) allows the release of the substrate IAA (magenta) to the extracellular side. NBD1 and NBD2 are in dark and light blue, while TMD1 and TMD2 are in dark and light orange.

