## Supplemental data for "The FKBP42, TWISTED DWARF1, prioritizes auxin over brassinosteroid transport by peptidyl-prolyl *cis-trans* isomerization of ABCB1"

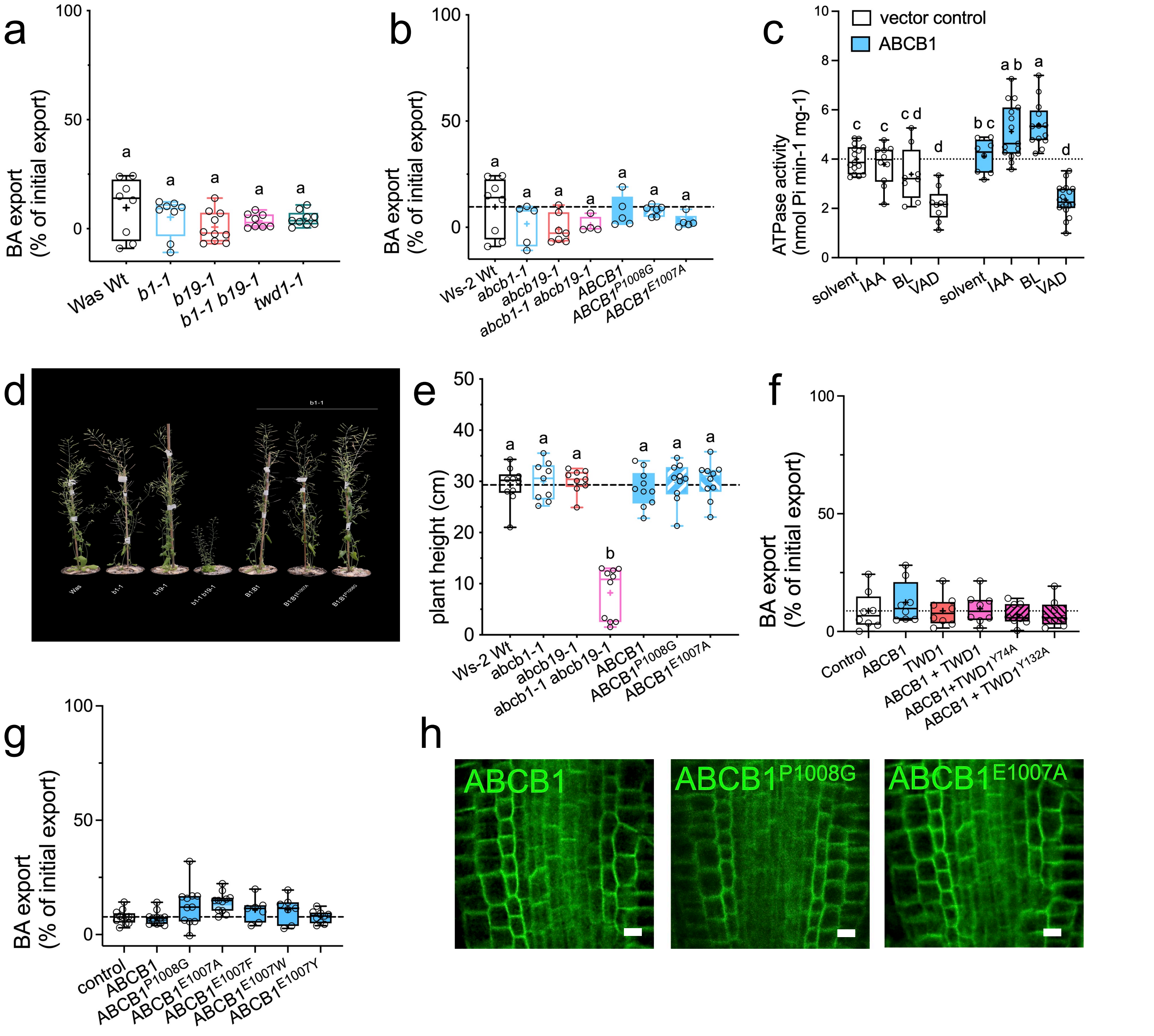


**Supplemental Fig. 1: Transport and *abcb1* complementation controls**

**a-b.** BA export from indicated Arabidopsis protoplasts. Significant differences ($p$ < 0.05) of means (indicated by "+") ± SE (n ≥ 8 independent protoplast preparations) were determined using Ordinary One-way ANOVA (Tukey's multiple comparison test) and are indicated by different lowercase letters.

**c.** ATPase activity of microsomal fractions prepared from tobacco leaves transfected with vector control or WT ABCB1 measured at pH 9.0 in the absence and presence of 5 µM IAA, BL or *ortho*-vanadate. Significant differences (p < 0.05) of means ± SE (n = 3 independent transfections and microsomal preparations) were determined using Two-way ANOVA and are indicated by different lowercase letters.

**d-e.** Complementation of *abcb1* with an IAA transport-incompetent version of ABCB1 ($ABCB1^{P1008G}$). Phenotype of 30 dag pot-grown plants (**c**) and quantification of plant height (**d**); bar, 5 cm.

**f-g.** BA export from indicated Arabidopsis (**e**) and tobacco (**f**) protoplasts. Significant differences ($p$ < 0.05) of means (indicated by "+") ± SE (n ≥ 8 independent protoplast preparations) were determined using Ordinary One-way ANOVA (Tukey's multiple comparison test) and are indicated by different lowercase letters.

**h.** Confocal imaging of Arabidopsis *abcb1-1* roots complemented with indicated mutant versions of ABCB1; bar, 50 µm.

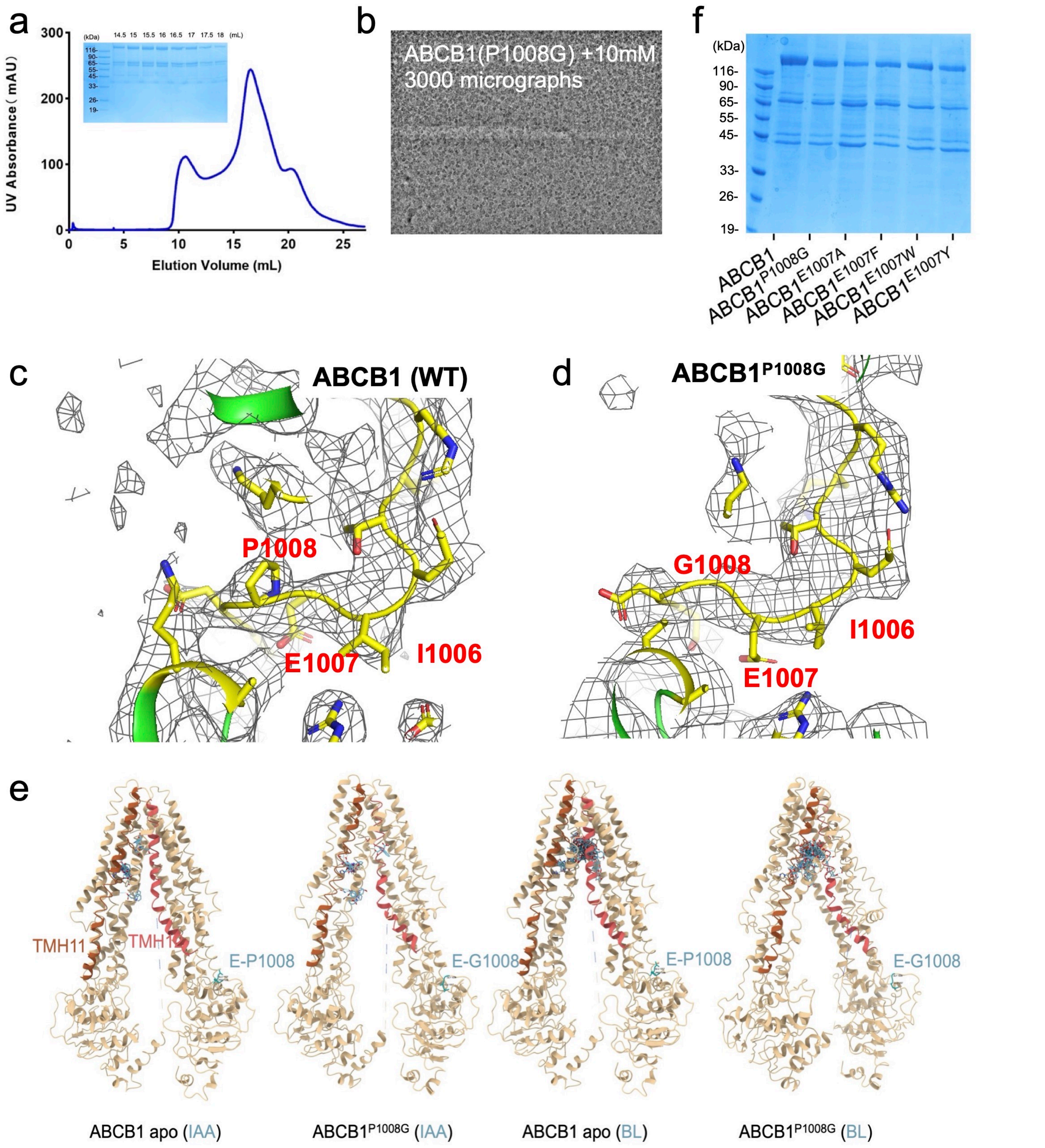


**Supplemental Fig. 2: ABCB1 expression and docking controls**

**a**. Representative gel filtration and Coomassie blue staining SDS-PAGE results (inset) of ABCB1$^{P1008G}$ co-expressed with TWD1 in HEK293F cells.

**b.** Typical cryo-EM image of the IAA-bound state of ABCB1$^{P1008G}$.

**c-d**. Cryo-EM structural comparison of the E-P1008 fold of Wt (apo) ABCB1 (**c**) and ABCB1$^{P1008G}$ (**d**) in IAA-bound states.

**e**. Front views of *in silico* docking (AutoDock Vina) of IAA and BL to Wt ABCB1 and ABCB1$^{P1008G}$ cryo-EM structures. TH11 and TH12 are colored in brown and red, respectively (see Fig. 2 for top views); IAA and BL in blue.

**f**. Representative Coomassie blue staining SDS-PAGE results of the wild-type (ABCB1) and E1007 mutants co-expressed with TWD1 in HEK293F cells after gel filtration.

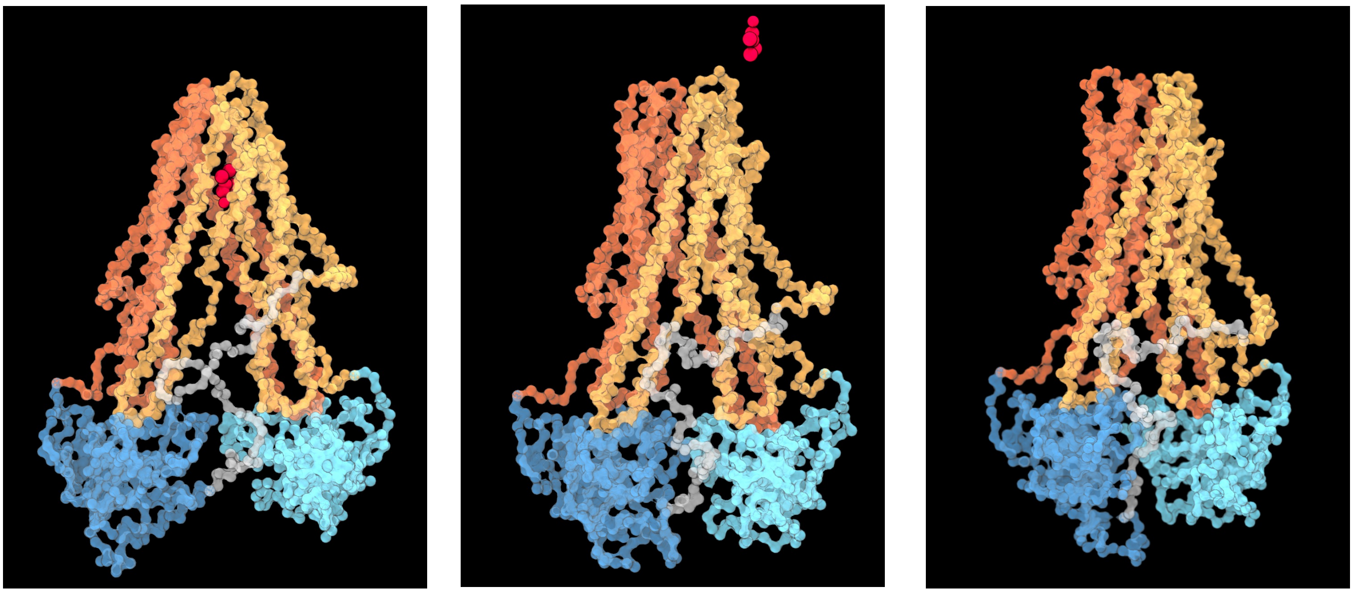

**Supplemental Fig. 3: CG-MD simulation controls**

ABCB1 conformational change from its open to its closed conformation, captured by CG-MD simulations.

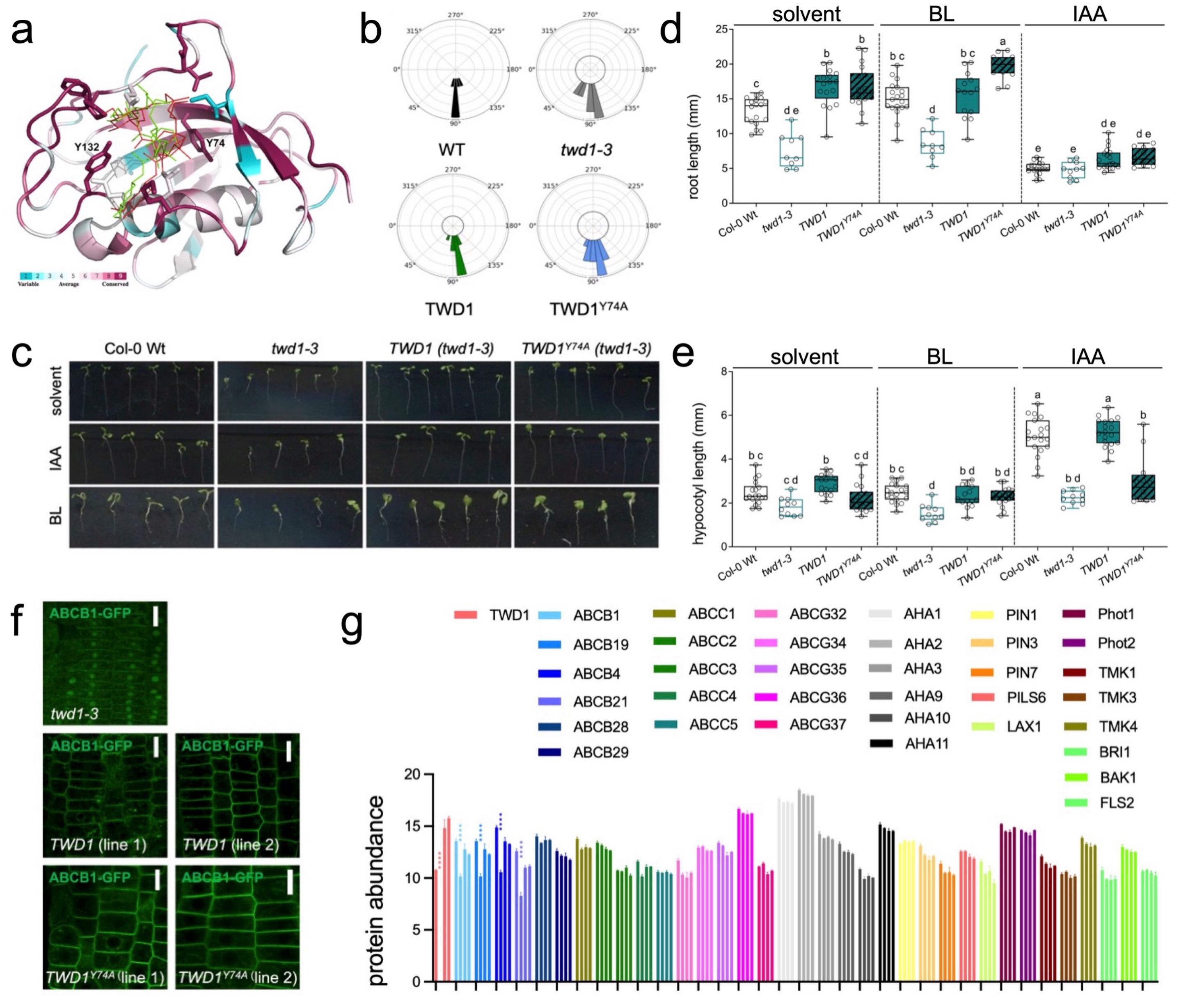

**Supplemental Fig. 4: *twd1* complementation controls**

**a.** Crystal structure (PDB 2F4E) of the TWD1 FKBD (aa 1-180) in complex with FK506 (green and red sticks) indicating the substrate binding pocket. Colors indicate Consurf conservation scores. Position of chosen FKBD mutations (Y74A, Y132A) facing toward the substrate and resulting in PPIase-dead versions of FKBP12 are indicated.

**b.** Helical wheel presentation of root gravitropism of *twd1-3* seedlings complemented with Wt TWD1 or TWD1$^{Y74A}$. Data show means of 3 experiments with each n ≥ 10 seedlings.

**c-e.** A PPIase-deficient (Y74A) version of *TWD1* complements growth defects of *twd1*. Phenotypes (**c**) and root (**d**) and hypocotyl length (**e**) of 7 dag seedlings grown on solvent control, IAA or BL (1 uM); bar, 1 cm. Significant differences ($p < 0.05$) of means (indicated by "+") to Wt solvent control ± SE (n ≥ 10 seedlings) were determined using Ordinary One-way ANOVA (Tukey's multiple comparison test) and are indicated by different lowercase letters.

**f.** Imaging of ABCB1-GFP in *ABCB1:ABCB1-GFP* (*twd1-3*) lines complemented with Wt *TWD1* or *TWD1$^{Y74A}$*, respectively; bar, 100 µm.

**g.** Mean abundance of ABCB1,4,19,21 and selected transporters and regulatory proteins relevant for this study. Microsomes of Wt (Col Wt, first column), *twd1-3* (second column) and *twd1-3* complemented with WT *35S:TWD1* or *35S:TWD1$^{Y74A}$* (two independent lines, columns three and four) were analyzed by label-free proteomics. Significant differences ($p < 0.05$) of means ± SE (n = 4 independent microsomal preparations and MS-MS analyses) to Wt control were determined using Two-way ANOVA followed by Tukey's multiple comparisons test are indicated by asterisks.

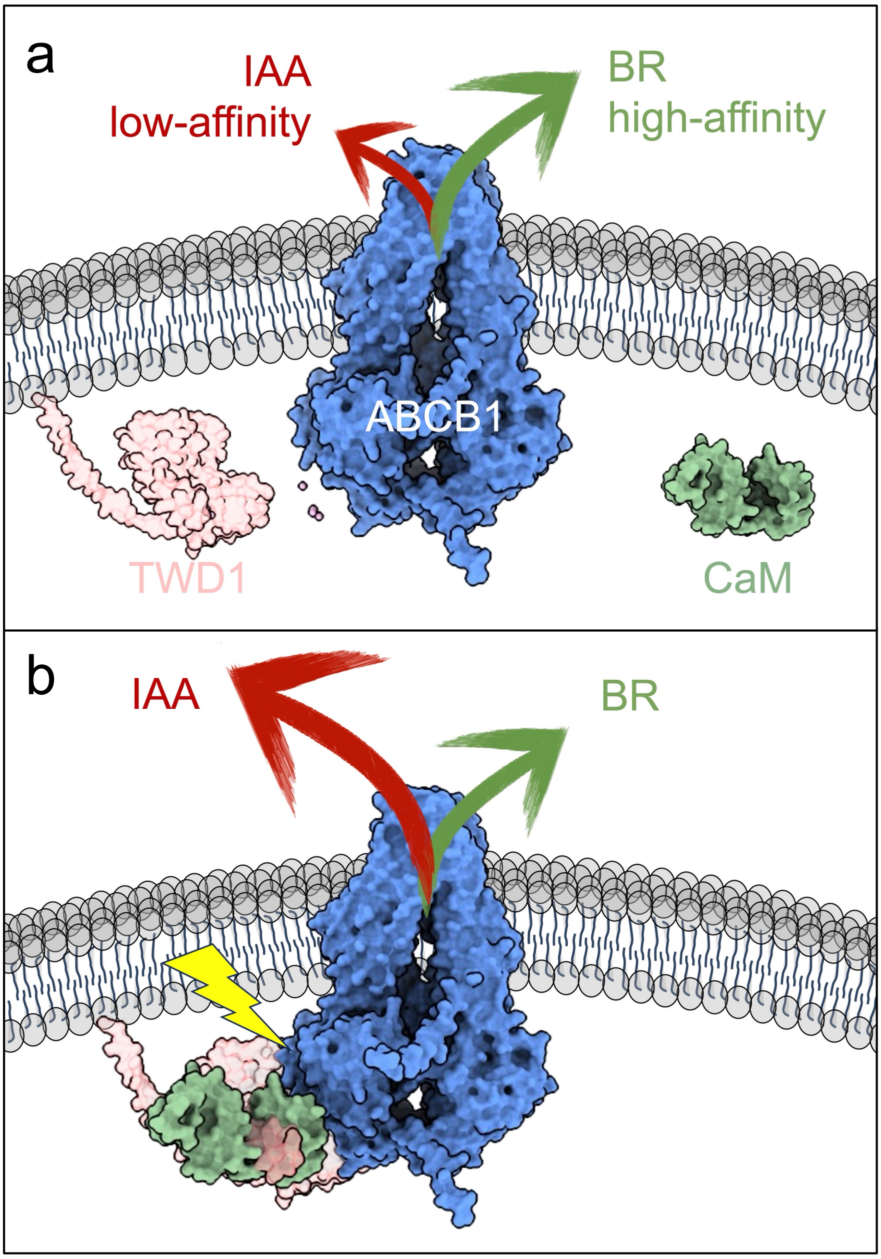


**Supplemental Fig. 5: Speculative model on unilateral IAA transport activation of ABCB1 by TWD1**

**a.** In a default state, ABCB1 would posses a low affinity for auxin (IAA) but high affinity for brassinosteroids (BR) allowing for both transport activities though with different capacities.

**b.** Recruitment of calmodulin (CaM), leads to an activation of a PPIase activity on TWD1, which is thought to confer ABCB1 into a high-affinity state for IAA by means of *cis-trans* peptidyl prolyl isomerization of a conserved D/E-P motif in the NBD2 of ABCB1 (yellow flash). Note that the calcium (not depicted here) might be essential for calmodulin activation but seems to be indispensable for PPIase activation.
